# Gain-of-function mutant p53 regulates long-noncoding RNA LINC00643 to modulate HIF1α in glioblastoma

**DOI:** 10.1101/2025.09.16.676559

**Authors:** Ying Zhang, Fang Yuan, Myron Keith Gibert, Collin Joseph Dube, Kadie Hudson, Cassandra Grello, Brian Reon, Shekhar Saha, Yunan Sun, Pawel Marcinkiewicz, Anindya Dutta, David Chief, Eric Holland, Roger Abounader

## Abstract

Gain-of-function mutations of p53 (GOF-MUT-p53) act as oncogenes by regulating gene transcription. We screened for the genome-wide transcriptional targets of GOF-MUT-p53 in glioblastoma (GBM) and found that a significant subset of them were long non-coding RNAs (lncRNA). Among these, LINC00643 was strongly repressed by GOF-MUT-p53 but not wild-type p53. LINC00643 was downregulated in GBM and low-grade glioma and correlated with patient survival. LINC00643 and its conserved third exon (Exon3) suppressed GBM cell proliferation, migration, invasion, stem cell self-renewal, and in vivo tumor growth Mechanistically, ChIRP-seq identified HIF1α as a key LINC00643 interactor. Under hypoxia, LINC00643 repressed HIF1α expression and its target genes by interacting with the HIF1α enhancer. Knockdown of GOF-MUT-p53 upregulated LINC00643 and reduces HIF1α, revealing a regulatory axis. These findings show extensive regulation of lncRNAs by GOF-MUT-p53 and uncover a novel mechanism by which GOF-MUT-p53 drives GBM through repression of LINC00643 and dysregulation of the HIF1α pathway.

## Introduction

Glioblastoma (GBM) is the most common and most deadly primary malignant brain tumor. GBM is characterized by rapid progression, resistance to standard therapies, and a dismal prognosis with an average life expectancy of ∼15 months ^1,2^. The Cancer Genome Atlas (TCGA) shows that the p53 pathway is deregulated in ∼85% of human GBM tumors ^3^. Upon closer analysis, the *TP53* gene was mutated in ∼28% of GBM ^3^. Most *TP53* mutations are missense point mutations that result in high expression of the mutant protein. The majority of these mutations occur within six “hotspots” between exons 4–8: R273, R248 R175, G245, R282, and R249 ^4^. Most of these mutant p53 (MUT-p53) are gain of function (GOF) that acquire oncogenic properties that actively drive tumor progression in addition to loss of the tumor-suppressive activities of wildtype p53 (WT-p53) ^5^. The modes of action of GOF-MUT-p53 are not fully understood but evidence suggests that one prominent mechanism is the regulation of transcription of a set of genes that are different from those regulated by WT-p53 ^6–8^.

Long non-coding RNAs (LncRNAs) are transcripts that do not have protein-coding potential and that are longer than 200 nucleotides. With flexible 3D molecular structure and dynamic cellular localization, lncRNAs regulate numerous physiological and pathological functions via a variety of mechanisms ^9,10,11,12^, many of which are still unknown. Among other, lncRNAs have been shown to act as protein scaffolds, chromatin regulators, microRNA sponges, regulators of mRNA stability and alternative splicing, and other mechanisms that regulate gene expression at the transcriptional, post-transcriptional, and epigenetic levels ^13,14,15,16,17,18^. LncRNAs have been reported to modulate neighboring transcription and participate in biological functions similar to those processed by nearby protein-coding genes ^19,20,21^. LncRNAs regulate critical biological functions, including cell growth, differentiation and development in normal and human disease, including GBM, through a variety of mechanisms ^22,23,24,25,26,27,28^. Recent studies report that lncRNAs play critical roles in cancer development by regulating several tumorigenic processes such as cell proliferation and apoptosis ^27,28,29^. However very few non-coding genes have been associated with GOF-MUT-p53 in GBM and cancer in general even though over 50,000 human lncRNAs have been identified ^9,10,11,12^ to date.

In this study, we screened for the transcriptional targets of GOF-MUT-p53 R273H in GBM using chromatin immunoprecipitation sequencing (ChIP-seq). We identified numerous GOF-MUT-p53 targets that differed from those of WT-p53. Surprisingly, among the 2,830 targets of GOF-MUT-p53 were 535 lncRNAs, the majority of which we also found enriched in MUT-p53 human tumors and deregulated in the TCGA database. This shows a previously underappreciated dimension of GOF-MUT-p53 functions and mechanisms of action. Furthermore, we found that the GOF-MUT-p53-regulated lncRNA LINC00643 is a tumor suppressor that acts by regulating HIF1α and hypoxia in GBM. Specifically, LINC00643 was downregulated in GBM and its expression is correlated with p53 mutational status and patient survival. LINC00643 ectopic expression inhibited GBM cell proliferation, migration, invasion, GBM stem cell (GSC) self-renewal and tumor growth in immunocompetent RCAS-Tva mice and immunodeficient xenografts. Mechanistically, LINC00643 bound to its neighbor gene HIF1α enhancer region and downregulated HIF1α expression and hypoxia signaling. We therefore show for the first time that GOF-MUT-p53 regulates a large subset of lncRNAs and uncover the GOF-MUT-p53-regulated LINC00643 as a new tumor suppressor that acts by regulating the expression of HIF1α and hypoxia in GBM.

## Results

### GOF-MUT-p53 binds to the promoters of numerous lncRNAs and regulates their expression in GBM

GOF-MUT-p53 has been shown to exert oncogenic effects, that are distinct from loss of WT-p53, by acquiring novel transcriptional activities, leading to the dysregulation of gene networks that promote tumor malignancy. We hypothesized that GOF-MUT-p53 drives aberrant transcriptional programs in GBM by regulating distinct sets of genes and pathways than WT-p53. To test this hypothesis, we performed ChIP-Seq in GOF-MUT-p53 and WT-p53 GBM cells. We used R273H (the most common p53 mutant in GBM) U373 cells and WT-p53 U87 cells (Fig. 1A). We identified 3406 unique binding sites for GOF-MUT-p53, denoted by peaks in binding activity relative to WT-p53, with less than 5% overlap between the two sets (Fig. 1B). This shows that R273H GOF-MUT-p53 has drastically different DNA binding loci than WT-p53. Among the most enriched peaks for GOF-MUT-p53 were 535 that were associated with lncRNAs loci and 2,120 that were associated with coding genes (Fig. 1B). Select GOF-MUT-p53 enriched lncRNAs are shown in Fig. 1C. To validate the GBM cell-based findings in human tumors, we analyzed the expression of these 535 associated lncRNAs in GBM tumors by correlating log10 fold changes in peak enrichment values from our ChIP-seq dataset with log2 gene deregulated genes in a R273H GOF-MUT-p53 GBM RNA-Seq dataset from the Cancer Genome Atlas (TCGA) and normal brain cortex datasets from the Genotype-Tissue Expression (GTEx) project (Fig. 1D). We found that 279 of the 535 lncRNAs significantly enriched in GOF-MUT-p53 were deregulated in MUT-p53 GBM patient tumors compared to normal brain cortex, with 109 downregulated and 170 upregulated lncRNAs (Figure 1D, blue and red dots, respectively). Thus, the majority of GOF-MUT-p53-regulated lncRNAs in GBM cells were also associated with the p53 mutational status in human GBM tumors. The binding of GOF-MUT-p53 to the promoters of select lncRNAs was verified by chromatin immune-precipitation (ChIP). DNA pulldown by p53 antibody was confirmed by qPCR, using known GOF-MUT-p53 target MYC as positive control and GAPDH as negative control (Fig. 1E, S1A). Altogether, the above data demonstrate that GOF-MUT-p53 regulates the expression of numerous lncRNAs in GBM cells and tumors.

**Figure 1.**
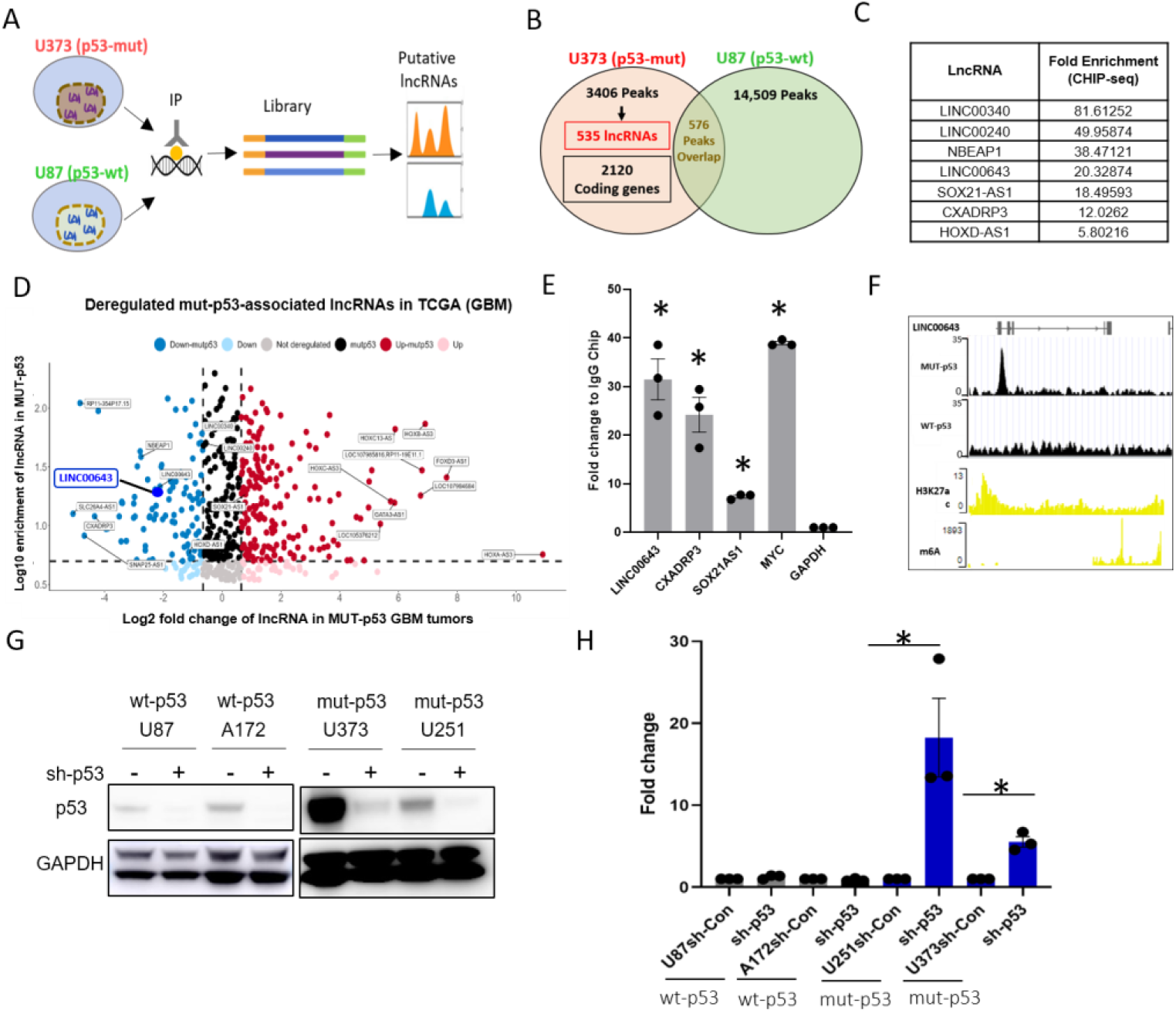
GOF-MUT-p53 directly regulates the expressions numerous lncRNAs in GBM. A) Schematics showing the screening strategy to identify GOF-MUT-p53 target genes in GBM cells. B) ChIP-seq peak analysis of GOF-MUT-p53 as compared with WT-p53. C) Selected lncRNA targets. D) Analysis of GOF-MUT-p53-bound lncRNAs for enrichment and deregulation in TCGA GBM database. E) ChIP validation of GOF-MUT-p53 binding to lncRNAs LINC00643, CXADRP3 and SOX21-AS1 using MYC as positive control and GAPDH as negative control. F) GOF-MUT-p53 binds to the LINC00643 promoter region (Genome Browser) compared to WT-p53 in GBM cells and H3K27a peak and m6A coverage in normal brain. G), H) p53 knockdown by shRNA in GBM cell line U87, A172 (with WT-p53) and U251, U373 (with MUT-p53 (R273H), validated by immunoblot in (G), showed that LINC00643 is upregulated after p53 knockdown in GOFMUT-p53 cells, but not in WT-p53 cells by qRT-PCR (H). *p= < 0.05

### Validation of GOF-MUT-p53 binding to lncRNA loci and regulation of lncRNA expression in GBM

We validated the binding of GOF-MUT-p53 to lncRNAs using ChIP in GBM cells expressing GOF-MUT-p53. The lncRNAs were LINC00643, SOX21-AS1 and CXADRP3 (Fig. 1C, 1D, S1A). They were selected based on their high deregulation (FC >= 1.6) in TCGA GBM RNA-Seq versus GTEx normal brain cortex samples and their enrichment for GOF-MUT-p53 binding (FC >= 10) in GBM Cell CHiP-Seq data. Among these, LINC00643 was prioritized for further study because of the absence of publications on it in GBM. A prominent GOF-MUT-p53 binding peak was identified upstream of the LINC00643 transcription start site (TSS), suggesting potential transcriptional regulation by this oncogenic transcription factor. To assess whether this region corresponds to an active promoter, we examined H3K27ac histone acetylation sites (for active regulatory regions) in the GTex Normal Tissue database. Consistent with active chromatin, a well-defined H3K27ac peak was detected in the same promoter-proximal region, supporting the notion that LINC00643 is transcriptionally active under physiological conditions (Fig. 1F). Additionally, analysis of m6A RNA methylation profiles (for RNA regulation or stability effects) in GTEx normal brain cortex samples revealed enrichment of m6A modifications at the 3′ end of LINC00643, implying post-transcriptional regulation through m6A-dependent mechanisms (Fig. 1F). Together, this analysis revealed histone acetylation at the LINC00643 promoter and m6A modification at its 3’ region suggesting that LINC00643 is under both transcriptional and epitranscriptomic control, and that its regulation may be perturbed in GBM through GOF-MUT-p53 binding. To determine if GOF-MUT-p53 regulates lncRNA expression, we knocked down GOF-MUT-p53 and WT-p53 with shRNA in the corresponding GBM cell lines. Knockdown efficiency was verified by immunoblotting (Fig. 1G). The data show that GOF-MUT-p53 knockdown leads to significant upregulation of LINC00643 expression measured by qRT-PCR, while WT-p53 knockdown did not lead to any significant change in LINC00643 expression (Fig. 1H). LncRNA SOX21-AS1 promoter and TSS enrichment by GOF-MUT-53 binding was shown in Fig. S1B and its expression was significantly downregulated by knocking down GOF-MUT-p53 (Fig. S1C). Together, these data show that GOF-MUT-p53 (R273H) binds to lncRNAs promoters and regulates their transcription.

### LINC00643 is downregulated in GBM and its expression correlates with GOF-MUT-p53 status and patient survival

Since GOF p53 mutations are common in GBM and because our data show that GOF-MUT-p53 inhibits LINC00643, we hypothesized that LINC00643 would be downregulated in GBM. Analysis of RNA-seq data of TCGA revealed that LINC00643 is significantly downregulated in GOF-MUT-p53 GBM (Fig. 2A, left) and MUT-p53 low grade glioma (LGG) (Fig. 2A, right) as compared to normal brain tissue from the GTEx Portal, and TCGA normal brain (Fig. S2C). Analysis of the Chinese Glioma Genome Atlas (CGGA) data showed that LINC00643 is significantly downregulated in IDH wildtype tumors compared to IDH mutant in both GBM and LGG (Fig. S2D). qRT-PCR analyses of banked human GBM specimens from the University of Virginia tumor bank confirmed that LINC00643 expression is significantly lower in tumors than in normal brain (Fig 2B). Furthermore, LINC00643 expression was notably lower in GOF-MUT-p53 GBM cell lines as compared to WT-p53 cell lines (Fig 2C). Importantly, analysis of survival data in TCGA and CGGA database showed that high expression of LINC00643 correlates with better survival outcomes in both GBM (Fig. 2D, left) and LGG patients (Fig. 2D, right). These data show that LINC00643 is downregulated in GOF-MUT-p53 gliomas and that downregulation associates with worse patient survival.

**Figure 2.**
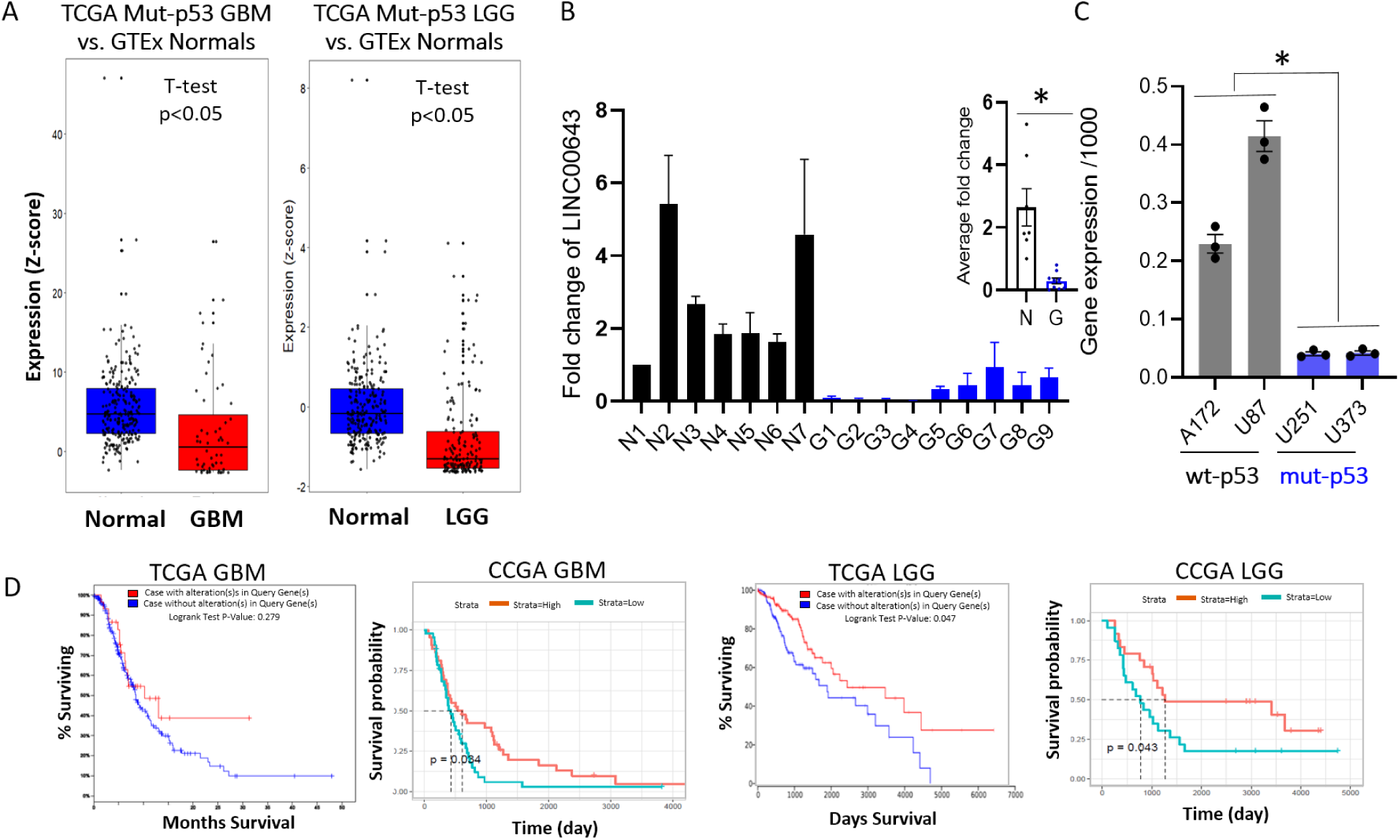
LINC00643 expression in downregulated in GBM tumors and in MUT-p53 GBM cells and is associated with patient survival. A) TCGA data analysis shows that LINC00643 is significantly downregulated in MUT-p53 GBM and low grade glioma (LGG). B) qRT-PCR validation of LINC00643 downregulation in banked GBM surgical specimens (G) vs normal brains (N) showing lower levels of expression as compared to normal brains with average fold change on right upper corner. C) LINC00643 expression in GBM cell lines (U251, U373) is downregulated as compared to WT-p53 (U87, A172) as measured by qRT-PCR. D) TCGA and Chinese Glioma Genomic Atlas (CGGA) survival analysis showing that high levels (red lines) of LINC00643 correlate with better survival in GBM (2 panels at left) and that high levels (red lines) of LINC00643 correlate with better survival in LGG (2 panels at right), respectively.

### LINC00643 inhibits GBM cell proliferation, migration, invasion, GSC self-renewal and GBM xenograft growth

To determine the functions of LINC00643 in GBM, we generated Tet-inducible stable cell lines to overexpress its full transcript in U251 GBM cells, G34 GBM stem cells (GSCs) (both express GOF-MUT-p53 and have low LINC00643 expression), and U87 WT-p53 cells. We assessed the effects of LINC00643 overexpression on cell growth, migration, invasion and GSC self-renewal. LINC00643 induction was confirmed by qRT-PCR following doxycycline (Dox) treatment (Fig. S3A). Overexpression of LINC00643 significantly inhibited GBM cell growth (Fig. 3A, left) and G34 self-renewal (Fig. 3A, right). Transwell invasion and cell scratch migration assays demonstrated that LINC00643 overexpression inhibits cell invasion (Fig. 3B) and migration (Fig. 3C). To investigate the effects of LINC00643 in vivo, we implanted the Tet-inducible LINC00643 GBM cells into the brains of immune-deficient mice. One week after tumor implantation, LINC00643 expression was induced by administering doxycycline to the animals. Tumors were visualized and their volumes measured by MRI three weeks later. The results showed a strong and significant inhibition of GBM growth following LINC00643 induction (Fig. 3D, S3B). These data collectively show that LINC00643 is tumor suppressive in GBM.

**Figure 3.**
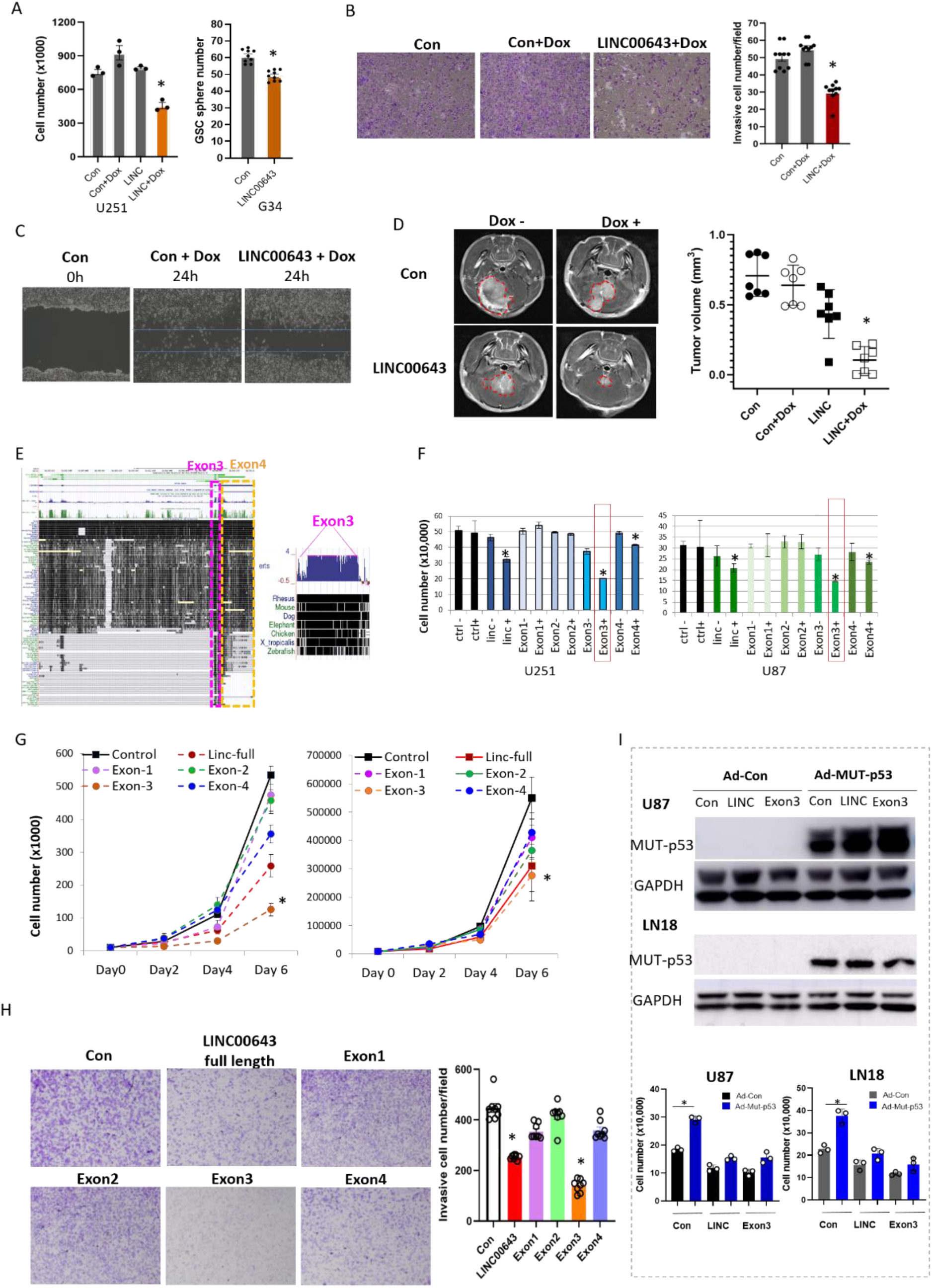
LINC00643 and conserved Exon3 inhibit GBM and GSC cell proliferation, migration, invasion and in vivo tumor growth. A) Cell growth and neurosphere assays showing that overexpression of LINC00643 by Tet-induction for 5 days in U251 cells and exogenous expression in GSCs G34 lead to a significant inhibition of cell growth and GSC self-renewal. B) Transwell invasion assay showing reduced invasion of U251 cell by Tet-induced LINC00643 (left) and quantification of invaded cells (right). C) Cell scratch assay showing that overexpression of LINC00643 by Tet-induction reduces migrating cells in the scratched area. D) GBM cells were intracranially implanted into the brains of immunodeficient mice (n=6). One week post tumor implantation, LINC00643 expression was induced by feeding the animals with doxycycline of 2 doxycycline + 5% sucrose for mg/mL). After 3 weeks, the mice were subjected to MRI imaging and tumor volume calculation. The results show that LINC00643 significantly inhibits in vivo GBM xenograft growth E) Genome browser analysis showing high conservation of LINC00643 Exon3 and partial conservation of Exon4 in multiple species, including human, mouse, rat, and chicken. F) Proliferation assay of individual exon-expressing GBM cell lines, U87 and U251 (left and right, respectively) showing that Tet-induced Exon 3 significantly inhibits cell proliferation in U251 (left) and U87 cells (right). G) Time course for cell proliferation assay showing consistent strong inhibitory effect of Exon3 in GBM cells U251 (left) and U87 (right). H) Invasion assay for different LINC00643 exons overexpression showing that Exon3 significantly inhibits GBM cell invasion compared to the full length LINC00643 and other exons (left) with quantified invasive cell numbers (right). I) LINC00643 and Exon3 mediate the oncogenic effects of GOF-MUT-p53. Immunoblot validation of overexpression of MUT-p53 in U87 or LN18 stable cells expressing LINC00643, Exon3 or control by transduction with adenovirus-control (Ad-Con) or Ad-MUT-p53 for 48 h (H upper panel). Proliferation assay shows that overexpression of LINC00643 or Exon3 reduced the oncogenic effect of MUT-p53 on GBM cell proliferation compared to control cells (H lower panel). * = p < 0.05.

### Exon 3 of LINC00643 is conserved and constitutes the functional part of the LINC00643

The LINC00643 gene consists of four exons, with Exon 3 being the most conserved based on a USCS genome browser and GETx analyses (Fig. 3E, S2E). To investigate the functional significance of the individual exons, we constructed plasmids for each of these exons, generated corresponding stable cells and confirmed their individual expressions (Fig. S3C). We then tested the effects of these exons on key malignant parameters in GBM cells, U251 and U87. We found that Exon 3 strongly inhibits GBM cell growth (Fig. 3F) and reduces cell proliferation (Fig. 3G). Additionally, Exon 3 significantly inhibited GBM cell invasion (Fig. 3H), with effects surpassing those of full-length LINC00643 (Fig. 3F, 3G, & 3H). Exons 1, 2 and 4 did not significantly alter the proliferation or invasion of GBM cells. These data indicate that Exon 3 is necessary and sufficient for the function of LINC00643.

### LINC00643 and Exon 3 mediate the effects of GOF-MUT-p53 in GBM

Having shown that GOF-MUT-p53 suppresses the expression of LINC00643 (Fig. 1H) and that LINC00643 is tumor suppressive in GBM, we hypothesized that suppression of LINC00643 mediates the oncogenic effects of GOF-MUT-p53. To test this hypothesis, we exogenously expressed GOF-MUT-p53 (R273H) via adenoviral transduction in U87 and LN18 GBM cells (which do not express GOF-MUT-p53) (Fig. 3I, upper panel). The cells were then Dox-induced to express LINC00643 or Exon 3, or control and the effects on cell growth were assessed. The data show that GOF-MUT-p53-induced GBM cell growth was significantly inhibited by overexpression of LINC00643 or Exon 3 (Fig. 3I, lower panel). These data show LINC00643 and Exon 3 mediate the effects of GOF-MUT-p53 in GBM.

### LINC00643 and Exon 3 inhibit GBM growth in RCAS-Tva and orthotropic xenograft mouse models and improve animal survival

We investigated the effects of LINC00643 on in vivo tumor growth and animal survival using an immunocompetent GBM mouse model and a GBM xenograft model. For the immunocompetent system, we utilized the RCAS-Tva-based GBM model (Fig. S4A), in which Nestin-Tva transgenic mice (*Ntv-a; Ink4a/Arf-/-; Pten flox/flox*) are intracranially injected with DF-1 cells producing RCAS-PDGFb and RCAS-Cre viruses. This model reliably induces high-grade gliomas with approximately 80% penetrance within 3– 5 weeks post-injection and recapitulates key features of human GBM^30^. To assess the role of LINC0043 in this context, we engineered RCAS vectors to express either the full-length or conserved exon 3. Because the human LINC00643 is ∼6.4 kb—exceeding the ∼3.5 kb capacity of the pRCAS vector—we constructed a truncated version containing Exons 1–3 and the conserved region of Exon 4 (pRCAS-hLinc) (Fig. S4B). We also generated RCAS constructs expressing only Exon 3 from human (pRCAS-hExon3) and mouse (pRCAS-mExon3), and the mouse homolog of LINC00643 (pRCAS-mLinc), or a scrambled control (pRCAS-Control) (Fig. S4B). DF-1 cells were transfected with these RCAS plasmids and injected intracranially into recipient mice alongside RCAS-PDGFb and RCAS-Cre DF-1 cells to initiate tumor formation. Tumor growth was monitored by MRI at 3–5 weeks post-injection. Expression of human LINC00643 and Exon3 led to a significant inhibition of spontaneous GBM tumor growth (Fig. 4A, S4D). LINC00643 and Exon3 also significantly improved the survival of the GBM bearing animals (Fig. 4B). Since Exon3 is highly conserved across species, we also performed similar experiments to the above using mouse homology of LINC00643 and Exon3. Expression of mouse LINC00643 and Exon3 similarly reduced GBM tumor growth compared to pRCAS-control mice (Fig. 4C, S4D) and prolonged animal survival (Fig. 4D), mirroring the effects observed with the human LINC00643 and Exon3. We also investigated the effects of LINC00643 and Exon3 in an immunodeficient mouse model of GBM. We implanted U251 and U87 stable cell lines expressing LINC00643 or Exon3 into the brains of immune deficient mice as well as control cells. Tumors were visualized and volumes were measured by MRI three weeks later. The data revealed strong and significant inhibition of in vivo GBM growth (Fig. 4E, 4G) with significantly prolonged animal survival (Fig. 4F, 4H). H&E staining of xenograft of LINC00643 and Control are shown in Fig. 4I and Fig. S4C. Immunofluorescent staining of tumor sections showed that the cell proliferation marker Ki67 is highly expressed in control tumors as compared to LINC00643 expressing tumors (Fig. 4J). These results demonstrate that LINC00643 and Exon3 have tumor-suppressive effects in GBM.

**Figure 4.**
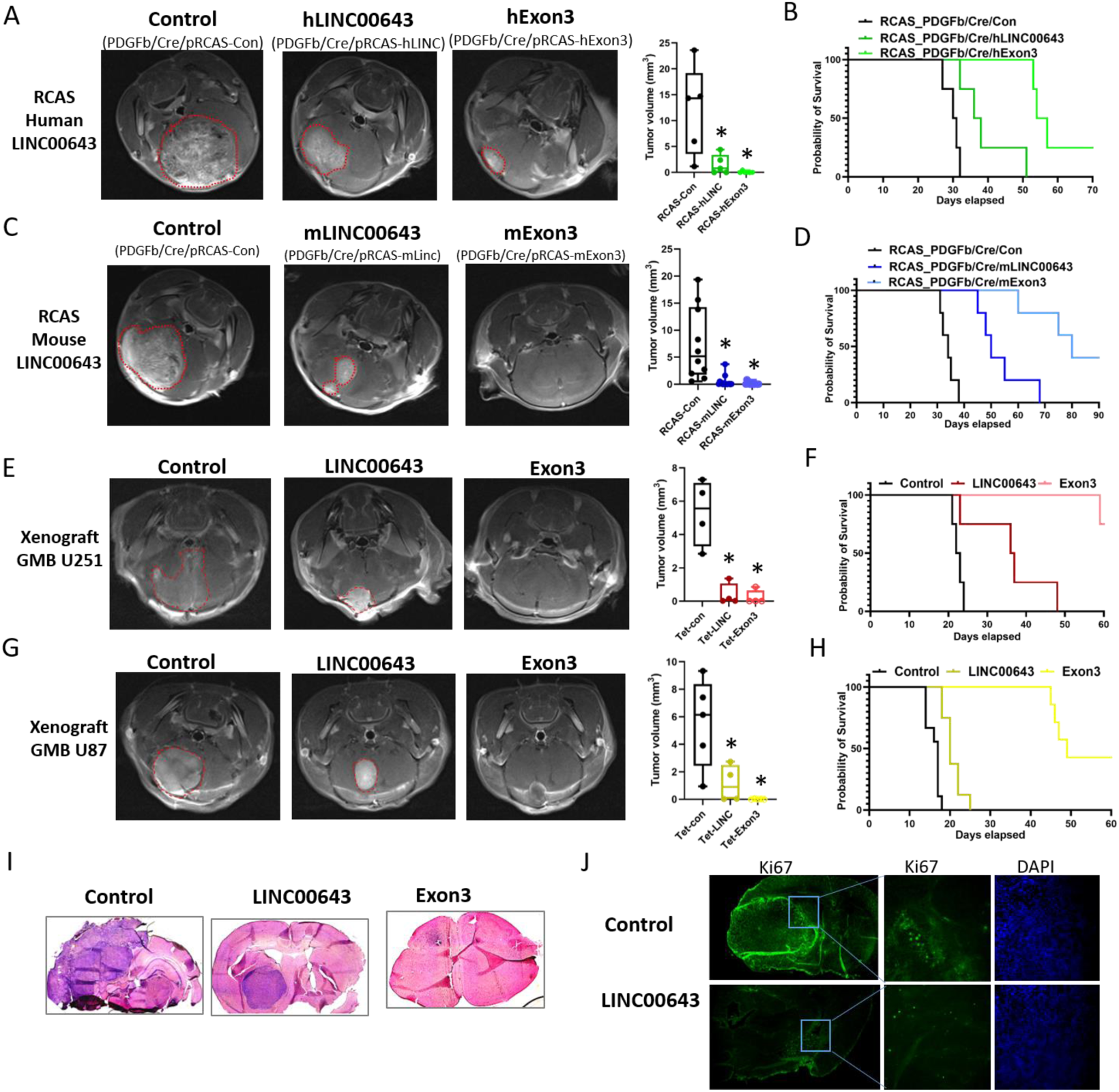
Human LINC00643 and Exon3 and mouse homologues significantly inhibit tumor growth in RCAS-Tva immune competent and in a xenograft GBM mouse model. A), B) We used the immune competent XFM/PDGFβ RCAS-Tva mouse to express LINC00643 or its highly conserved Exon3 in GBM tumors. DF-1 cells were co-transfected with pRCAS-PDGFb/Cre + p-RCAS-Control or p-RCAS-LINC00643/Exon3 for 12 days and were intracranial injected in the brain of RCAS-Tva mice N/tv-a; Ink4a-arf-/-; Ptenfl/fl. The animals were subjected to MRI imaging 4 weeks after injection and animal survival was assessed. Expression of both human LINC00643 and Exon3 led to a strong and significant inhibition of GBM tumor growth (A) and significantly improved animal survival (B). C), D) Expression of mouse homologues of LINC00643 and Exon3 also resulted in significant reduction of tumor growth (C) and significant better survival (D). E), F) U251 GBM cells expressing Tet-inducible human LINC00643 and Exon3 were implanted in the brains of immune deficient mice. One-week post tumor implantation, LINC00643 or Exon3 expression were induced by feeding the animals doxycycline. Tumors were visualized and their volumes measured by MRI three weeks later. The results show a strong and significant inhibition of in vivo GBM growth after LINC00643 or Exon3 expression (E). LINC00643 and Exon 3 significantly extended the survival of tumor bearing animals (F). G), H) U87 cells with Tet-inducible human LINC00643 / Exon3 were xenografted in immunodeficiency mice that were fed doxycycline one week after tumor implantation. A significant inhibition of GBM tumor growth was observed (G) with better survival (H). I) H&E staining of RCAS-Tva GBM tumors that overexpress LINC00643, Exon3 and control. J) IHC staining of Ki67 of RCAS tumor showed that overexpression of LINC00643 results in reduced Ki67 expression compared to control tumors. * = p < 0.05.

### LINC00643 localizes to the nucleus

The cellular localization of lncRNAs is important for their mechanisms of action. To determine the localization of LINC00643, we performed single molecule RNA fluorescent in situ hybridization (smRNA FISH). In GOF-MUT-p53 U251 control cells, LINC00643 expression was not detectable, consistent with the low levels of LINC0064 in Fig. 2C. However, in LINC00643-overexperssing stable cells U251-LINC00643, LINC00643 was predominantly localized in the nucleus (Fig. 5A). This finding was further confirmed by cell fractionation followed by qRT-PCR using U6B and U44 as nucleic controls and GAPDH and PPIA as cytoplasmic controls, which demonstrated that LINC00643 is primarily nuclear (Fig. 5B). Nuclear lncRNAs can act as key regulators of gene expression through diverse mechanisms such as chromatin remodeling, transcriptional regulation, and epigenetic modification. They interact with chromatin modifiers, transcription factors, or DNA elements to influence gene expression at the transcriptional level.

**Figure 5.**
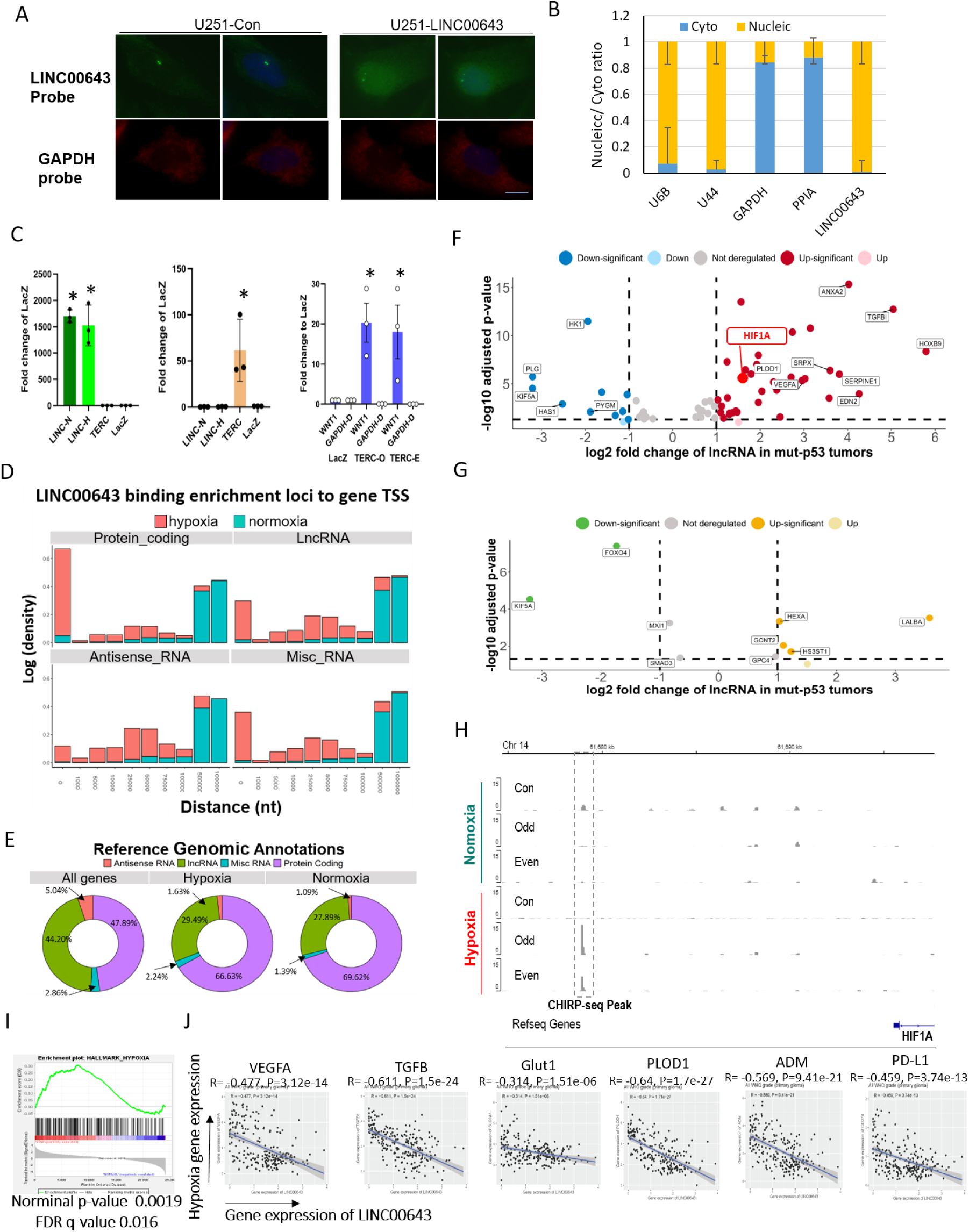
LINC00643 regulates HIF1α transcription by binding to the HIF1α 5’ enhancer region. A) B) LINC00643 is mainly located in nucleus as shown by by smRNA FISH (A) and cell fractionation followed by qRT-PCR (B). C) (Left panel) ChIRP was performed using U251-LINC00643 stable cell lysate and probe sets of LINC probe set, or LacZ probe set (negative control), TERC probe set (positive control), respectively. Retrieved RNA was then analyzed by qRT-PCR. LINC00643 were detected in LINC probes pulldown cell lysis, but not in TERC and GAPDH samples, suggesting specificity of LINC00643 pulldown. (Middle panel) TERC was detected in TERC probe samples, but not Laz-probe while unspecific GAPDH and LINC was not detected in TERC and Lac probe samples, confirmed the specific TERC probe pull down. (Right panel) WNT1 promoter is known bound by TERC which was used as positive control. WNT1 was detected by qPCR using pulled down DNA with TERC-O and TERC-E probes, but not by LacZ probe. D) The ChIRP-seq result analysis of LINC00643 binding enrichment distance to the transcription start site (TSS) of protein coding genes, lncRNAs, Antisense RNAs and MiscRNAs. LINC00643 binds to closer regions to TSS under hypoxia (Red) compared to normoxia (Blue). E) The ChIRP-seq reference genomic Annotations at hypoxia and normoxia growth conditions. F) Volcano plot showing that many hypoxia hallmark, hypoxia response, and angiogenesis genes that associate with LINC00643 under hypoxic conditions are significantly deregulated. Red genes are significantly upregulated, while blue genes are significantly downregulated G) Volcano plot showing that fewer hypoxia hallmark, hypoxia response, and angiogenesis genes that associate with LINC00643 under normoxic conditions are significantly deregulated. Gold genes are upregulated, while green genes are downregulated H) ChIRP-seq analysis showing that LINC00643 binds to 5’ of HIF1α region suggesting that it might regulate HIF1α expression as co-factor for HIF1α transcription. I) Gene set enrichment assay of illumina Hiseq (n=101) the cancer genome atlas (TCGA) showed significant enrichment of gene sets involved in the hypoxia with the abundant genes being upregulated (cutoff: median). J) Correlation expression of LINC00643 with hypoxia-associated genes in CGGA database. * = p < 0.05.

### LINC00643 is located near the HIF1a locus

Many nuclear lncRNAs are often involved in regulating nearby gene transcription. Notably, the LINC00643 genomic locus is close to that of the alpha unit of hypoxia inducible factor 1 (HIF1α). This a key gene in human embryonic development that is also heavily involved in cancer and GBM progression, specifically through direct involvement in the regulation of genes under hypoxic conditions. In addition to being co-located to the HIF1α locus, the tumor-suppressive effects of LINC00643, specifically Exon 3, are much stronger in GBM mouse models than in cells in vitro. Since the gene is operating under hypoxic conditions *in vivo* and normoxic conditions *in vitro*, we therefore hypothesized that LINC00643 may interact with HIF1α, leading to differential effects under hypoxic and normoxic conditions.

### LINC00643 binds different genomic targets under hypoxic and normoxic conditions

To identify whether LINC00643 has differential effects under hypoxic and normoxic conditions, we first sought to identify which genes interact with LIN00643 in these contexts. Many of these genes will have known functions and identifying which ones bind with LINC00643 can therefore provide insight into LINC00643’s function. Some may even be directly related to hypoxia and HIF1α and would therefore provide critical insight into the relationship between HIF1α and LINC00643. To accomplish this goal, we performed Chromatin Isolation by RNA purification (ChIRP) followed by deep sequencing (ChIRP-seq) to identify LINC00643 DNA binding partners (Fig. S5A). This enabled us to map its genome-wide chromatin interaction landscape under hypoxic and normoxic conditions. We sought to determine if LINC00643 binds to genomic DNA in GBM cells under normoxic conditions. We used the known DNA-binding lncRNA TERC as positive control and LacZ, a non-human gene, as a negative control^31^. Using qRT-PCR of RNAs pulled down by probes targeting LINC00643, TERC, or LacZ, we confirmed the specific retrieval of LINC00643 (Fig. 5C, left) and TERC (Fig. 5C, middle). Importantly, we observed significant enrichment of WNT1 promoter regions, which are known TERC binding sites, by qPCR of pulled down DNAs using TERC probes, validating our positive control (Fig. 5C, right). This shows that, in our control experiments, LINC00643 can bind to genomic DNA under normoxic conditions, a critical control for ensuing high throughput experiments.

After confirming that LINC00643 can bind genomic DNA, we sought to identify all genomic DNA sites where LINC00643 binds. To accomplish this, we divided the LINC00643 specific probes into two groups (odd and even) based on their 5’ to 3’ sequence order and analyzed the binding peaks observed in both groups (Fig. S5B, S5C), with each binding peak representing unique binding sites in the genome. We identified 38347 genomic regions that LINC00643 binds under hypoxic conditions and 5720 under normoxic conditions, with 3556 overlaps (Fig. S5D). The ChIRP-seq annotation of the probes bound to these sites revealed that LINC00643 binds to several sites located near the transcription start sites (TSS) of both protein-coding and non-coding genes (Fig. 5D), though there was not an appreciable difference in the types (only the quantity) of unique genes that interact with LINC00643 under hypoxic or normoxic conditions (Fig. 5E). These data suggests that, while LINC00643 binds similar types of genes under hypoxic and normoxic conditions, it is far more transcriptionally active under hypoxia than normoxia, binding many more unique genes.

We then integrated the LINC00643 ChIRP-Seq enrichment data, showing LINC00643-associated genes, with TCGA GBM RNA-Seq data, using methodologies previously described, to identify which LINC00643-associated genes under hypoxic conditions were also deregulated in GBM Tumors. From 38347 probes that bound to LINC00643 within 20kb of the TSS, we identified 6145 statistically significant (FDR <= 0.05) upregulated (FC >=2) and 4603 downregulated (FC <= -2) genes in GBM Tumors. When focused on linc00643-associated genes that are hallmark hypoxia, hypoxia response, and angiogenesis genes, we found that several were deregulated under hypoxic conditions (Fig. 5F). Fewer of these genes were deregulated under normoxic conditions (Fig. 5G). Critically, these deregulated linc00643-associated genes under hypoxic conditions included hypoxia-related protein coding genes such as the critical HIF1α, along with VEGFA, HOXA3, MYBL2, TOP2A, CEP55, PPP5D1, NSF, and FAM126B, and lncRNAs such as HOXA-AS3,

RP1170O19,22 and RP1170O19.23. (Fig. 5F, S5D, S5E). Conversely, we found far fewer LINC00643-associated genes under normoxic conditions (n = 3556) that were also statistically significantly upregulated (n = 586) or downregulated (n = 595) in GBM Tumors (Fig. 5G). Notably, we were able to experimentally demonstrate that LINC00643 significantly binds to the 5’ UTR and coding region of the neighboring gene HIF1α (Fig. 5H), suggesting that LINC00643 may directly regulate HIF1α transcription, leading to its deregulation under hypoxic conditions. Interestingly, the binding region is located at -16301 nt upstream from the HIF1α transcription start site (TSS), a region that has been identified as a conserved enhancer region of HIF1α in human craniofacial development ^32^. To further understand the underlying mechanism by which LINC00643 regulate HIF1α, we performed Gene Set Enrichment Analysis (GSEA) using TCGA GBM patient data. We found that low expression of LINC00643 was associated with upregulation of several hypoxia-related genes, including those involved in hypoxia regulation and angiogenesis (Norminal p-value 0.0019, FDR q-value 0.016) (Fig. 5I, S5F). We also found that LINC00643 expression is inversely correlated with hypoxia gene expression in TCGA and CGGA database, including VEGFA, TGFB1, GLUT1, PLOD1, HOXB9, ADM, PD-L1, and several others (Fig. 5J, S5G), many of which are significantly deregulated under hypoxic conditions as previously mentioned.

### LINC00643 and Exon3 inhibit HIF1α expression

Based on the proximity of the LINC00643 to the HIF1α gene locus, on the ChIRP-seq enrichment of LINC00643 at the HIF1α 5’ enhancer, and on the correlation between LINC00643 expression and hypoxia pathways in TCGA and CGGA databases, we hypothesized that LINC00643 regulates HIF1α transcription via an RNA-dependent mechanism. To test this hypothesis, we overexpressed LINC00643 and Exon3 in GBM cells grown under hypoxic conditions for various time points, then examined HIF1α expression using qRT-PCR and Immunoblotting, comparing the results to control cells grown under normoxic conditions. The data showed that overexpression of both LINC00643 and Exon3 resulted in significant downregulation of HIF1α mRNA at early time points (10min, 30min and 1hr) (Fig. 6A) and HIF1α protein level at later time points (8 hrs and 24 hrs) (Fig. 6B) in U251 and U87 cells.

**Figure 6.**
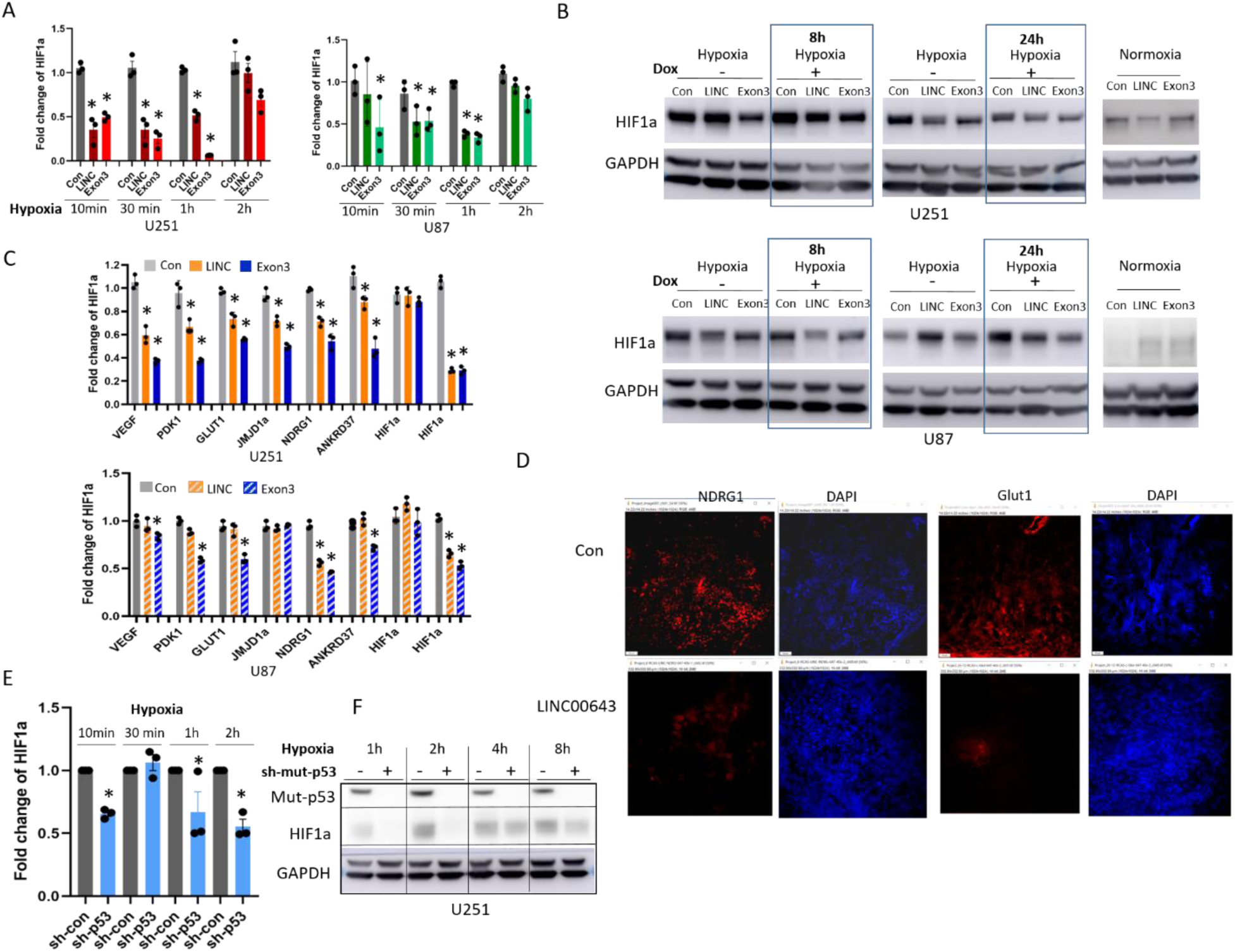
GOF-MUT-p53 regulates HIF1α expression under hypoxic conditions via suppression of LINC00643. A) Tet-inducible LINC00643 and Exon3 GBM cells were treated with or without doxycycline (Dox) for 24 h and grown under hypoxia condition for different time points and assessed for HIF1α RNA expression by qRT-PCR. The results show that LINC00643 and Exon3 inhibit HIF1α transcriptional expression at 10 min, 30 min and 1 h. B) Dox-inducible LINC00643 and Exon cells were grown under hypoxia for 8 hrs and 24 hrs and HIF1 α protein levels were determined by immunoblotting and compared to normoxia condition. The data show that LINC0063 and Exon3 induction reduced HIF1 α protein level at 8 hrs and 24 hrs. C) LINC00643 and Exon3 overexpression downregulated hypoxia-regulated gene expression, including VEGFA, PDK1, GLUT1, NDRG1, and HIF2α under hypoxia as measured by qRT-PCR. D) Representative immunofluorescent staining of NDRG1 and Glut1 (Red) in Control and LINC00643 overexpressing mouse tumors showing that overexpression of LINC00643 reduces NDRG1 and Glut1 expression. E) F) MUT-p53 was knocked down with shRNA in U251 GBM cells grown under hypoxic conditions for different times and assessed for HIF1 α RNA by qRT-PCR (E) and protein by immunoblotting (F). MUT-p53 knockdown was validated by immunoblotting (F). *\* = P < 0.05*

Next, we assessed the expression of several known hypoxia-associated genes using qRT-PCR after overexpression of LINC00643 and Exon3. We observed that the RNA level of VEGFA, PDK1, GLUT1, JMJD1a, NDRG1, ANKRD37 and HIF2α were significantly reduced upon overexpression of LINC00643 and/or Exon3 followed by hypoxia incubation for 48 hrs (Fig. 6C). We also assessed the hypoxia associated protein expression in mouse xenografts with overexpression of LINC00643 or control by immuno-florescent staining of tumor section. We found that NDRG1 and GLU1 expression are significantly inhibited in LINC00643-overexpressing tumors compared to control tumors (Fig. 6D). NDRG1 and GLUT1 are highly relevant hypoxia-responsive genes in GBM and known as key indicators of how tumors adapt to low-oxygen environments ^33,34^. This shows that LINC00643 regulates hypoxia pathways in GBM.

GOF-MUT-p53 has not been reported to regulate HIF1α RNA and protein expression via lncRNA in GBM, although one study has suggested that a protein-protein interaction of GOF-MUT-p53 and HIF1α regulates hypoxia transcription in gastric and esophageal adenocarcinoma cells ^31^. Since GOF-MUT-p53 directly binds the LINC00643 promoter and represses its expression, we investigated whether GOF-MUT-p53 regulates HIF1α expression at transcription or protein level in GBM cells. In sh-GOF-MUT-p53 knockdown GBM cells grown under hypoxic conditions, HIF1α RNA expression was significantly reduced at 10 min, 1hr and 2hrs (Fig. 6E). Additionally, HIF1α protein expression was significantly inhibited at multiple time points (1hr, 2hrs, 3hrs and 4hrs) (Fig. 6F, S6B). These data show that GOF-MUT-p53 regulates HIF1α expression by regulating LINC00643.

In summary, our studies shows for the first time that GOF-MUT-p53 regulates a subset of lncRNAs in GBM. Among these lncRNAs, GOF-MUT-p53 binds to the gene and suppresses the expression LINC00643 leading to its downregulation in GBM and LGG, where LINC00643 low expression correlates with poor patient survival. Functionally, LINC00643 suppresses tumor growth and GSC stemness. Mechanistically, LINC00643 suppresses HIF1α transcription and hypoxia signaling by binding to the HIF1α enhancer. These findings reveal a novel GOF-MUT-p53-lncRNA-HIF1α axis that regulates hypoxia and malignancy in GBM.

## Discussion

GBM is the most aggressive brain cancer with very few good therapeutic options. The frequently altered TP53 gene in GBM often sustains missense mutations in the DNA-binding domain. Several of these mutations are known to confer gain-of-function properties to the highly expressed mutant protein in many cancers ^35–39^. Among these mutations, GOF-MUT-p53 (R273H) is one of the most frequent mutations in GBM, where it promotes cell proliferation, invasive growth, GSC stemness through only partially understood mechanisms. One likely mechanism is the regulation of the transcription of sets of genes that differ from those regulated by WT-p53. To uncover GOF-MUT-p53 mechanisms of action, we designed a p53 ChIP-seq strategy in endogenous GOF-MUT-p53 and WT-p53 GBM cells to identify transcriptional targets specific to hotspot missense GOF-MUT-p53. Surprisingly, and for the first time to our knowledge, we found that a substantial fraction (535) of GOF-MUT-p53-bound genes were lncRNAs. The association between GOF-MUT-p53 and lncRNAs was confirmed in human tumor data sets. We then conducted deep functional and mechanistic studies on lncRNA LINC00643 in the context of the GOF-MUT-p53-driven GBM progression. We found that LINC00643 expression is significantly downregulated in GBM and LGG and its higher expression is correlated with better survival in the TCGA and CGGA database. Functionally, we showed that upregulation of LINC00643 impairs GBM cell proliferation, migration and invasion. LINC00643 also acted as a tumor suppressor in an immunocompetent RCAS-Tva and an immunodeficient xenograft mouse model of GBM. Rescue experiments showed that LINC00643 downregulation mediated the oncogenic effects of GOF-MUT-p53 (R273H). Mechanistically, we showed that the GOF-MUT-p53-LINC00643 axis acts via regulation of HIF1 α expression.

Our genomic analysis showed that Exon3 of LINC00643 is highly conserved across species (Fig. 3E). Consistently, we observed that Exon3 exhibits a strong tumor-suppressive effect both in vitro and in vivo, suggesting that Exon3 might be the functional domain of LINC00643. LINC00643 and Exon3 showed significant stronger inhibitory effects on primary tumor growth in mouse models compared to in vitro experiments. These results suggest that LINC00643’s tumor-suppressive effects may be particularly relevant to specific growth condition and microenvironment found in vivo. Previous studies have reported a requirement for MUT-p53 in mouse models of T cell lymphomas^40^ and non–small cell lung cancer ^41^, where tumor cells exhibit a dependency/addiction on the gain-of-function activity of MUT-p53. In GBM, tumor cells may become similarly dependent on GOF activity of MUT-p53 for maintenance and survival. Overexpression of LINC00643 or Exon3 could block GFO-MUT-p53 signaling and disrupt its oncogene addiction and thereby repress tumor growth.

We found that GOF-MUT-p53 binds to the promoter region of LINC00643 and inhibits its transcription. LINC00643 is predominantly localized in nucleus, and its genomic locus is situated near HIF1α. LncRNAs have been implicated in genome scaffolding and gene regulation of nearby genes. Using ChIRP-seq analysis, we found that LINC00643 binds to 5’ enhancer region of HIF1α, leading to the downregulation of HIF1α RNA and protein level under hypoxic conditions. We therefore propose a model according to which LINC00643 binds to the enhancer of the HIF1α gene and prevents enhancer promoter interaction to inhibit HIF1α transcription. By inhibiting LINC00643 expression, GOF-MUT-p53 reverses this process leading to HIF1α activation (Figure 7). Consistent with this model, we observe downregulation of HIF1α when GOF-MUT-p53 was knocked down by shRNA.

**Figure 7.**
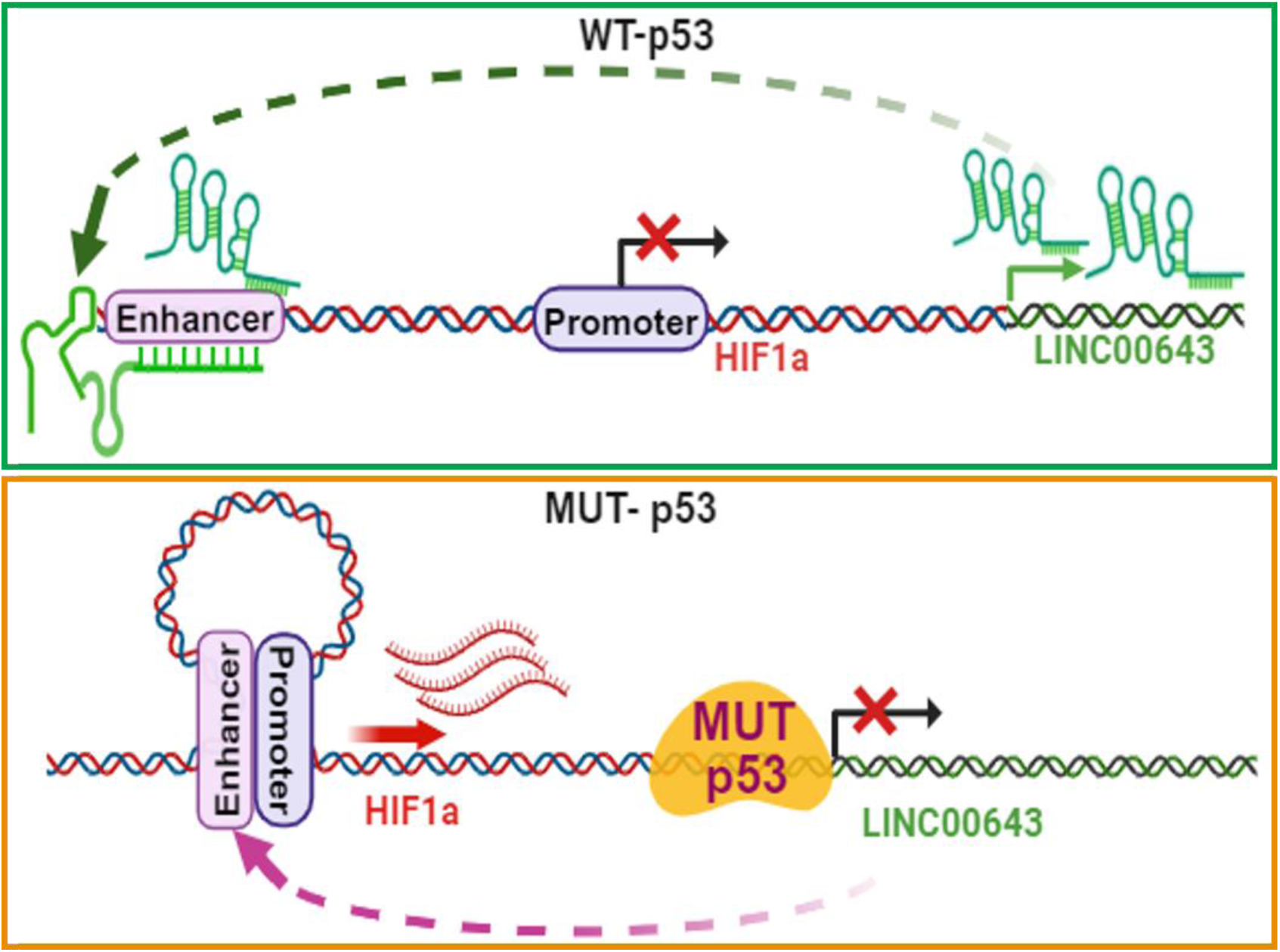
Schematic summary. In WT-p53 cells, LINC00643 is highly expressed and binds to the HIF1α enhancer region to repress its transcription, thereby suppressing hypoxia signaling. In contrast, in GBM or LGG cells harboring GOF-MUT-p53, the mutant protein binds to the LINC00643 promoter and suppresses its expression. As a result, LINC00643 can no longer occupy the HIF1α enhancer, allowing enhancer–promoter interaction and activation of HIF1α transcription. This leads to increased HIF1α expression and induction of hypoxia-related pathways.

One of hallmarks of GBM progression is the hypoxic environment in the tumor that enables cancer cells to utilize HIF-dependent transcriptional regulation to promote malignancy ^42^. HIF1α activates the transcription of numerous genes involved in angiogenesis, stemness, invasion, autophagy, and metabolic reprogramming ^43–46^, all of which contribute to advanced tumor properties in hypoxic conditions. Under normal oxygen conditions, HIF1 α is rapidly degraded by the oxygen-dependent degradation domain (ODDD) through the VHL-mediated proteasomal pathway ^47^. However, HIF1α protein remains stable under hypoxia. We show that this process is partly regulated by GOF-MUT-p53 and LINC00643. Our findings suggest a new model in which the LINC00643 directly represses HIF1α transcription by binding to an enhancer region associated with the HIF1α locus. This repression occurs rapidly (within 10 minutes to 1 hour), preceding the observed decrease in HIF1α protein levels at later time points (8–48 hours). This temporal pattern suggests that the lncRNA does not regulate HIF1α via post-translational mechanisms such as protein degradation or destabilization, which are well-characterized modes of HIF1α regulation under normoxic conditions. Instead, our data support a transcriptional mechanism, whereby lncRNA binding may interfere with enhancer-promoter looping, recruit transcriptional repressors or chromatin modifiers, or alter chromatin accessibility at the HIF1α locus. Notably, the subsequent downregulation of canonical hypoxia-responsive genes at 48 hours further supports a model in which lncRNA suppresses the HIF1α transcriptional program through early inhibition of HIF1α transcription initiation or enhancer activity. This represents a previously unrecognized layer of regulation within the hypoxia response pathway, mediated by lncRNA-directed transcriptional control.

While there is expansive literature on the link between WT-p53 and HIF1α / hypoxia ^48^, very few studies have explored the association between GOF-MUT-p53 and HIF1α in cancers. Recent studies have investigated the connection between mutant p53 and HIF1α to promote hypoxia signaling in gastric and esophageal adenocarcinoma cell lines ^49^. In non–small cell lung cancer, a physical interaction between GOF-MUT-p53 and HIF1α has been shown to regulate extracellular matrix components to promote cancer progression ^50^. GOF-MUT-p53 has also been implicated in increasing tumor vascularization via ROS-mediated activation of the HIF1α / VEGFA pathway in colon tumor cells ^51^. We show for the first time that GOF-MUT-p53-regulated LINC00643 attenuates HIF1α in GBM.

Although LINC00643 downregulation is significantly correlated with missense p53 mutation in GBM, some GBM specimens with WT-p53 exhibit relatively low expression compared to normal brain tissue. This suggests that GOF-MUT-p53 may not be the sole regulator of LINC00643. The brain consumes approximately 20% of total body’s oxygen, despite representing only 2% of total body weight ^52^. Prolonged hypoxia in the brain leads to neuronal cell death via apoptosis, contributing to hypoxic brain damage ^53,54^. LINC00643, especially Exon1-3, is highly expressed in various regions of normal brain compared to other tissues (Fig. S2B, S2E), while HIF1α expression in normal brain is significantly low (Fig. S5H), indicating their association. In this study, we demonstrated that LINC00643 represses HIF1α expression suggesting it may play an important role in normal brain function and maintenance and tumor progression. This suggests that multiple molecules and mechanisms, beyond GOF-MUT-p53, could be involved in the regulation of LINC00643. Further investigation into LINC00643 and its association with hypoxia signaling could improve our understanding of GBM and inspire new therapeutic strategies.

In summary, our study uncovers a previously unrecognized regulatory axis in GBM, wherein GOF-MUT-p53 (R273H) directly regulates lncRNAs, identified as a transcriptional targets uniquely enriched in GOF-MUT-p53 GBM cells. Among the lncRNAs, LINC00643 is significantly downregulated in GBM and LGG, and its expression correlates positively with patient survival. We demonstrate that LINC00643—particularly its conserved third exon—acts as a potent tumor suppressor by inhibiting GBM cell proliferation, invasion, stemness, and in vivo tumor growth in both immunocompetent and xenograft models. Mechanistically, LINC00643 binds to the HIF1α enhancer and suppresses its transcription, leading to reduced HIF1α protein levels and downstream hypoxia signaling. This repression occurs independently of protein stabilization mechanisms and precedes functional inhibition of the hypoxia response. These findings define a novel LINC00643-mediated mechanism by which GOF-MUT-p53 promotes gliomagenesis, highlighting LINC00643 as a potential biomarker and therapeutic agent to counteract GOF-MUT-p53 oncogenic pathways.

## Materials and Methods

### Ethical approval

The ethics committee of the University of Virginia approved data access. The methods used in this work were conducted in accordance with the approved guidelines. The studies were approved by an institutional review board. Further information and requests for raw data and resources should be directed and will be fulfilled by the Contact: Roger Abounader. All animal experiments were approved by the Institutional Animal Care and Use Committee at University of Virginia.

### Data Availability Statement

Raw fastq reads and processed peak files for GOF-MUT-p53 ChIP-Seq in U87 cells, WT-p53 CHIP-Seq in U373 cells and LINC00643 ChIRP-Seq (under hypoxic and normoxic conditions) in U87 cells, data generated for this project, are provided via a publicly-accessible Gene Expression Omnibus (GEO) project: (insert GEO code here). Detailed methodologies and processed data are provided as supplement to this document (codebook.zip) and online at <u>github.com/abounaderlab/gof_mutp53_lncrna_project</u>. Please refer to README.md document (<u>github.com/abounaderlab/gof_mutp53_lncrna_project</u>) for additional instructions for downloading and extracting repositories, code, and supplementary data, and the corresponding author for any data access questions.

### Detailed Computational Methodologies - Codebook

Detailed methods, including additional information for all datasets used, can be found as a codebook provided in the supplementary materials and online at: <u>github.com/abounaderlab/gof_mutp53_lncrna_project</u>. This repository contains the processed data and codes to regenerate certain manuscript figures, with a Markdown document for each figure with section headers. The UNIX command line (bash shell) and/or R programming languages were used for all analyses contained in the codebook. Please refer to README.md document on <u>github.com/abounaderlab/gof_mutp53_lncrna_project</u> for additional instructions for downloading and extracting repositories, code, and supplementary data, and the corresponding author for any data access questions.

### Cell culture and specimens

GBM Cells lines U87, A172, U373, and other used cells 293T and DF1 were purchased from American Type Culture Collection (Manassas, VA). U251MG was a gift from Dr. Benjamin Purow. GBM stem cell G34 (a kind gift from Dr. Jeongwu Lee, Cleveland Clinic) were isolated from patient surgical specimens and characterized for *in vivo* tumorigenesis, pluripotency, self-renewal, stem-cell markers, and neurosphere formation. Normal human astrocytes (NHA) were purchased from Lonza Bioscience. Cell lines that were used for more than 6 months after purchase were reauthenticated by STR profiling. GBM surgical specimens and normal brain were obtained from the University of Virginia Brain Tumor Bank according to procedures that were reviewed and approved by the Review Board of the University of Virginia Health System. U87 cells were cultured and maintained in Minimal Essential Medium (MEM) supplemented with one mM sodium pyruvate, one mM non-essential amino acids (NEAA), 10 ml of 7.5% sodium bicarbonate, 1% penicillin/streptomycin, and 10% fetal bovine serum. U373 cells were cultured in DMEM low glucose medium with 1% penicillin/streptomycin and 10% fetal bovine serum. U251 cells were grown in RPMI medium with 1% penicillin/streptomycin and 10% fetal bovine serum. A172 and 293T cells were cultured in DMEM high glucose medium supplemented with 1% penicillin/streptomycin and 10% fetal bovine serum. Glioblastoma stem cells, G34 cells were cultured in Neuron Basel Medium (500ml) consisting 2.5 ml of N-2 (0.5X), 5 ml of B27 (0.5X), 50 ng/ml EGF, 50 ng/ml FGF, 0.5 mM glutamine, and 1% penicillin/streptomycin. The NHA cells were cultured in growth medium kit from Lonza Bioscience (Durham, NC, USA). All cells were cultured at 37°C in a humidified incubator with a 5% CO2 atmosphere to maintain cell viability.

### TCGA AND GTEx RNA-Seq Data

GBM (n = 161) and LGG (n = 504) RNA-Seq data were acquired from the Cancer Genome Atlas via the dbGAP portal and were compared to normal brain cortex from TCGA and the Genotype-Tissue Expression Database (GTEx, n = 260) acquired via the ANVIL portal. Bowtie2 was used to align raw reads to the hg38 genome. Bedtools were used to convert files to sam, bam, and bed formats. R/Rstudio was used for all downstream analyses. Detailed methods, including additional information for all datasets used, can be found as a codebook provided in the supplementary materials and online at: <u>github.com/abounaderlab/gof_mutp53_lncrna_project</u>.

### Differential expression and Survival Analyses using TCGA Datasets

TUCR expression, deregulation, and survival association, were analyzed using processed TCGA and GTEx RNA-Seq data and a workflow using R/RStudio. TPM scores were generated to assess absolute expression. DESeq2 and z-scores were used to calculate deregulated genes. The cox coefficient of proportional hazards (CPH) from survminer and Kaplan-Meier plots acquired via oncolnc.org were used to assess correlation with patient outcomes. Detailed methods, including additional information for all datasets used, can be found as a codebook provided in the supplementary materials and online at: <u>github.com/abounaderlab/gof_mutp53_lncrna_project</u>.

### Chromatin immunoprecipitation and deep sequencing (ChIP-seq)

To identify target genes that are regulated by GOF-mut-p53, but not wt-p53 we performed ChIP-seq on GOF-MUT-p53 U373 cells and WT-p53 U87 cells. Chromatin was crosslinked to fix the protein-DNA complexes with 1% formaldehyde for 10 minutes at room temperature, followed by quenching with 125 mM glycine for 5 minutes at room temperature. Cells were harvested and lysed in a buffer containing protease inhibitors. Chromatin was sheared into fragments of approximately 200–500 base pairs using a Branson sonicator. Immunoprecipitation was carried out using a p53 antibody that recognizes both mutant and wild-type p53 protein (Cell Signaling, Danvers, MA), followed by incubation with protein A/G magnetic beads. The protein-DNA complexes were washed, eluted, and crosslinking was reversed by incubating at 65°C overnight. DNA was then purified using a spin column and prepared for sequencing. The library preparation and sequencing were conducted by (ZymoResearch, Tustin, CA).

### Single-molecule fluorescence in situ hybridization (smFISH)

To determine LINC00643 localization, LINC02273 RNA FISH was performed using Stellaris FISH probes as listed in Supplementary Table S2 (Biosearch Technologies, California USA) according to the manufacturer’s protocol. In brief, U251-LINC00643 and U251-Control stable cell lines were grown on coverslips in a 24-well culture plate. Cells were fixed with 4% (w/v) paraformaldehyde in 1xPBS for 10min and permeabilized with 0.5% Triton X-100. After washing with PBS, cells were incubated with LINC00643 or GAPDH specific probe sets that were labeled with (Quasar 570 green for LINC00643 and CAL Fluro 590 red for GAPDH) to allow for the detection of individual RNA molecules. The hybridization step was performed at an optimal temperature of 37C overnight to ensure probe binding. Following hybridization, unbound probes were washed away, and the samples were mounted with a mounting medium containing DAPI (VECTASHIELD Antifade Mounting Medium, Vector Laboratories, Newark, CA USA) to stain nuclei. Fluorescence microscopy was used to capture images of the cells, allowing for the subcellular visualization LINC00643.

### Chromatin isolation by RNA purification and deep sequencing (ChIRP-seq)

Chromatin isolation by RNA purification (ChIRP) was performed to analyze LINC00643-chromatin interactions. LINC00643 probes were designed using LGC Biosearch’s web-based designer, biotin-TEG-labeled at its 3′ terminus and divided into odd or even groups (Suppl Methods and Materials). Probes targeting LacZ were used as negative control. The GOF-MUT-p53 U251-Tet-LINC cells were treated with Dox (10 ng/µl) for 24 hours and continue incubated at hypoxia (1% Oxygen) or normoxia condition (20% Oxygen) for another 24 hr. For each sample, U251 cells (4 × 10^7^) were crosslinked with 1% glutaraldehyde (Sigma, St. Louis, MO) for 10 min at room temperature, followed by quenching with 125 mM glycine for 5 min. The cells were then harvested, washed and lysed using a Branson sonicator in ChIRP lysis buffer containing RNase inhibitors and sheared to 100–500 bp fragments at 4 °C. Cleared lysates were incubated with biotinylated probes to hybridize to the chromatin-RNA complexes in 37 °C for overnight. After hybridization, the RNA-chromatin complexes were captured using streptavidin-coated magnetic beads. The beads were washed extensively to remove non-specific binding, and the RNA and DNA were purified accordingly. The isolated RNA and DNA were then purified using standard extraction methods. qPCR and RT-qPCR were performed as described above. Library construction for pulled down DNA samples were used for ChIRP libraries preparation using the DNA SMART™ ChIP-Seq Kit for Illumina (Takara, San Jose, CA) according to the manufacturer’s instructions. The libraries were sent to UVA Genome Analysis Technology Core for Agilent sample analysis and Illumina Next Gen Sequencing.

### Processing of raw ChIP- and ChIRP-Seq

Raw fastq output reads were aligned by Bowtie2 (v2.2.9) with a human reference genome (hg38). The uniquely mapped reads were subjected to the peak calling algorithm, MACS (v1.4.2) with default parameters. Genes were considered enriched if there was at least a 10-fold increase in GOF-MUT-p53 binding versus WT-p53 (ChIP-seq), or for LINC00643 binding versus input (ChIRP-seq). Detailed methods, including additional information for all datasets used, can be found as a codebook provided in the supplementary materials and online at: <u>github.com/abounaderlab/gof_mutp53_lncrna_project</u>.

### Integration of ChIRP-Seq results with processed TCGA datasets

The expression and deregulation (via log2FC) in GBM (and LGG) of 42,644 genes in our dataset were calculated using previously described methods. Raw ChIRP-Seq fastq reads were converted to peak files (with enrichment values) using previously described methods. Bedtools (v1.10) closest was used to identify the nearest genomic element. The tidyverse RStudio package was used to integrate these two datasets, using gene aliases as the joining factor. For ChIRP-Seq, enriched (FC >= 10) genes that were deregulated by at least 2-fold (log2FC >=1, FDR <= 0.05) in TCGA GBM Tumors versus normal brain cortex were considered differentially expressed. These analyses were performed on cells expressing LINC00643 that were grown under either hypoxic or normoxic conditions. Detailed methods, including additional information for all datasets used, can be found as a codebook provided in the supplementary materials and online at: <u>github.com/abounaderlab/gof_mutp53_lncrna_project</u>.

### Animal experiments

Animal experiments were approved by the University of Virginia Animal Care and Use Committee (ACUC) (Protocol Number: 3542-05-21). The immunodeficient Hsd:Athymic Nude-Foxn1 (Nude) mice were from Inotiv (Indianapolis, IN, USA). The RCAS-Tva mice (*Ntv-a Ink4a-Arf-/-LPTEN)* were bred. Female and male mice were used equally in this study.

Orthotopic xenograft formation: To generate human GBM xenografts, mice were anesthetized with ketamine (100 mg/kg body weight), and placed on a sterotactic frame. U251 or U87 cell lines (3X10^5^) that express either Dox-inducible LINC00643, Exon3 or control were intracranially implanted in the right striata at the coordinates from the bregma 1 mm anterior, 1.5 mm lateral and 2.5 mm intraparenchymal. After tumor cell implantation, mice were removed from the sterotactic apparatus, kept in separate cages, checked for signs and symptoms of neurologic deficits and then euthanized if positive. Three weeks after tumor implantation, the animals were subjected to brain MRI (7 Tesla Bruker/Siemens ClinScan small animal MRI). To measure the tumor size, 20 µl of gadopentetate dimeglumine (Magnevist, Bayer Healthcare, Whippany, NJ) was intraperitoneally injected 15 minutes before scanning. Tumor volumes were measured using MicroDicom. For survival studies, mice were monitored carefully and euthanized when they displayed symptoms of tumor development.

RCAS-Tva mouse model: To assess LINC00643 effects on spontaneous GBM tumor formation in the RCAS-Tva mouse model, DF-1 cells were transfected with plasmids of RCAS-Scrambled, RCAS-hLINC00643, RCAS-hExon3, RCAS-mLINC00643, RCAS-mExon3, RCAS-PDGFB, or RCAS-Cre. Mice used for this study were anesthetized with ketamine, xylazine as mentioned above and placed on a stereotactic frame. Two microliters (1:1:1 mixtures of RCAS-PDGFB, RCAS-CRE and either RCAS-Scrambled, RCAS-hLINC00643, RCAS-hExon3, RCAS-mLINC00643, or RCAS-mExon3) of 4 x 10^4^ transfected DF-1 cell suspensions were injected into the subventricular zone with the stereotactic coordinates 1.0 mm lateral to midline, 1.0 mm anterior to bregma, 2.5mm deep to the cranial surface. Mouse brains were scanned with MRI at 6-8 weeks after cell implantation. Tumor volumes were measured using MicroDicom and survival studies were conducted as described above.

### Statistics

All experiments were repeated at least three times (except microarrays that were conducted twice in triplicates). Two group comparisons were analyzed with a *t*-test, *P*-values were calculated, and these values were adjusted for multiple hypothesis correction using the Bonferroni method. All quantitative results are shown as mean ± standard error of the mean (SEM). Molecular experiment tests were performed using Sigmaplot 14, and computational experiment tests were performed using RStudio. *FDR*<=0.05 was considered as significant and symbolized by an asterisk in the figures.

## Conflict of Interest

The authors have declared that no conflicts of interest exist

## Funding

Supported by NIH/NINDS RO1 NS122222, NIH/NCI UO1 CA220841, NIH/NINDS 1R21NS122136, NCI Cancer Center Support Grant P30CA044579, a University of Virginia Comprehensive Cancer Center Pilot Grant, a Schiff Foundation grant (all to R.A.). Support was also provided by the University of Virginia Advanced Microscopy Facility, Molecular Imaging Core, and Genome Analysis and Technology Core.

## Authorship

Designing research studies: YZ and RA. Conducting experiments and acquiring data: YZ, FY, MG, CD, KH, CG, BR, SS, YS, PM. Analyzing Data: YZ, MG, KH, RA. Conceptual input: YZ, RA, AD, DC, EH. Writing manuscript: YZ and RA. Reviewing and editing of manuscript: All authors

## Notes

### Competing Interest Statement

The authors have declared no competing interest.

## References

1 . Wen, P. Y. & Reardon, D. A. Neuro-oncology in 2015: Progress in glioma diagnosis, classification and treatment. Nat. Rev. Neurol. 12, 69–70 (2016).

2 . Wen, P. Y. & Kesari, S. Malignant gliomas in adults. N. Engl. J. Med. 359, 492–507 (2008).

3 . Brennan, C. W. et al. The somatic genomic landscape of glioblastoma. Cell 155, 462–477 (2013).

4 . Walker, D. R. et al. Evolutionary conservation and somatic mutation hotspot maps of p53: Correlation with p53 protein structural and functional features. Oncogene 18, 211–218 (1999).

5 . Dittmer, D. et al. Gain of function mutations in p53. Nat. Genet. 4, 42–46 (1993).

6 . Peart, M. J. & Prives, C. Mutant p53 gain of function: the NF-Y connection. Cancer Cell 10, 173–174 (2006).

7 . Zalcenstein, A. et al. Mutant p53 gain of function: repression of D95 (Fas/APO-1) gene expression by tumor-associated p53 mutants. Oncogene 22, 5667–5676 (2003).

8 . Xu, J. et al. Gain of function of mutant p53 by coaggregation with multiple tumor suppressors. Nat. Chem. Biol. 7, 285–295 (2011).

9 . Fatica, A. & Bozzoni, I. Long non-coding RNAs: new players in cell differentiation and development. Nat. Rev. Genet. 15, 7–21 (2014).

10 . Mercer, T. R. & Mattick, J. S. Structure and function of long noncoding RNAs in epigenetic regulation. Nat. Struct. Mol. Biol. 20, 300–307 (2013). 10.1038/nsmb.2480

11 . Heo, J. B., Lee, Y. S. & Sung, S. Epigenetic regulation by long noncoding RNAs in plants. Chromosome Res. 21, 685–693 (2013). 10.1007/s10577-013-9392-6

12 . Ramos, A. D., Attenello, F. J. & Lim, D. A. Uncovering the roles of long noncoding RNAs in neural development and glioma progression. Neurosci. Lett. 625, 70–79 (2016). 10.1016/j.neulet.2015.12.025

13 . Wang, K. C. et al. A long noncoding RNA maintains active chromatin to coordinate homeotic gene expression. Nature 472, 120–124 (2011).

14 . Engreitz, J. M. et al. The Xist lncRNA exploits three-dimensional genome architecture to spread across the X chromosome. Science 341, 1237973 (2013).

15 . Kretz, M. et al. Control of somatic tissue differentiation by the long non-coding RNA TINCR. Nature 493, 231–235 (2013).

16 . Anderson, K. M. et al. Transcription of the non-coding RNA upperhand controls Hand2 expression and heart development. Nature 539, 433–436 (2016).

17 . Engreitz, J. M. et al. Local regulation of gene expression by lncRNA promoters, transcription and splicing. Nature 539, 452–455 (2016).

18 . Paralkar, V. R. et al. Unlinking an lncRNA from its associated cis element. Mol. Cell 62, 104–110 (2016).

19 . Engreitz, J. M. et al. Local regulation of gene expression by lncRNA promoters, transcription and splicing. Nature 539, 452–455 (2016).

20 . Joung, J. et al. Genome-scale activation screen identifies a lncRNA locus regulating a gene neighbourhood. Nature 548, 343–346 (2017).

21 . Olivero, C. E. et al. p53 activates the long noncoding RNA Pvt1b to inhibit Myc and suppress tumorigenesis. Mol. Cell 77, 761–774 (2020).

22 . Rinn, J. L. & Chang, H. Y. Genome regulation by long noncoding RNAs. Annu. Rev. Biochem. 81, 145–166 (2012).

23 . Batista, P. J. & Chang, H. Y. Long noncoding RNAs: Cellular address codes in development and disease. Cell 152, 1298–1307 (2013).

24 . Mercer, T. R. & Mattick, J. S. Structure and function of long noncoding RNAs in epigenetic regulation. Nat. Struct. Mol. Biol. 20, 300–307 (2013).

25 . Heo, J. B., Lee, Y. S. & Sung, S. Epigenetic regulation by long noncoding RNAs in plants. Chromosome Res. 21, 685–693 (2013).

26 . Ramos, A. D., Attenello, F. J. & Lim, D. A. Uncovering the roles of long noncoding RNAs in neural development and glioma progression. Neurosci. Lett. 625, 70–79 (2016).

27 . Qi, P. & Du, X. The long non-coding RNAs, a new cancer diagnostic and therapeutic gold mine. Mod. Pathol. 26, 155–165 (2013).

28 . Spizzo, R., Almeida, M. I., Colombatti, A. & Calin, G. A. Long non-coding RNAs and cancer: A new frontier of translational research? Oncogene 31, 4577–4587 (2012).

29 . Wahlestedt, C. Targeting long non-coding RNA to therapeutically upregulate gene expression. Nat. Rev. Drug Discov. 12, 433–446 (2013).

30 . Hambardzumyan, D. et al. Modeling adult gliomas using RCAS/t-va technology. Transl. Oncol. 2, 89–98 (2009).

31 . Chu, C., Qu, K., Zhong, F. L., Artandi, S. E. & Chang, H. Y. Genomic maps of long noncoding RNA occupancy reveal principles of RNA–chromatin interactions. Mol. Cell 44, 667–678 (2011).

32 . Wilderman, A., VanOudenhove, J., Kron, J., Noonan, J. P. & Cotney, J. High-resolution epigenomic atlas of human embryonic craniofacial development. Cell Rep. 23, 1581–1597 (2018).

33 . Said, H. M. et al. Oxygen-dependent regulation of NDRG1 in human glioblastoma cells in vitro and in vivo. Oncol. Rep. 21, 237–246 (2009).

34 . Nishioka, T. et al. Distribution of the glucose transporters in human brain tumors. Cancer Res. 52, 3972– 3979 (1992).

35 . Dong, P. et al. Mutant p53 gain-of-function induces epithelial–mesenchymal transition through modulation of the miR-130b-ZEB1 axis. Oncogene 32, 3286–3295 (2013).

36 . Freed-Pastor, W. A. et al. Mutant p53 disrupts mammary tissue architecture via the mevalonate pathway. Cell 148, 244–258 (2012).

37 . Freed-Pastor, W. A. & Prives, C. Mutant p53: one name, many proteins. Genes Dev. 26, 1268–1286 (2012).

38 . Kogan-Sakin, I. et al. Mutant p53(R175H) upregulates Twist1 expression and promotes epithelial– mesenchymal transition in immortalized prostate cells. Cell Death Differ. 18, 271–281 (2011).

39 Schofield, H. K., et al. Mutant p53R270H drives altered metabolism and increased invasion in pancreatic ductal adenocarcinoma. JCI Insight 3, e97422 (2018).

40 . Alexandrova, E. M. et al. Improving survival by exploiting tumour dependence on stabilized mutant p53 for treatment. Nature 523, 352–356 (2015).

41 . Vaughan, C. A. et al. Addiction of lung cancer cells to GOF p53 is promoted by up-regulation of epidermal growth factor receptor through multiple contacts with p53 transactivation domain and promoter. Oncotarget 7, 12426–12446 (2016).

42 Kaelin, W. G., Jr. How oxygen makes its presence felt. Genes Dev. 16, 1441–1445 (2002).

43 . Lee, J.-W., Bae, S.-H., Jeong, J.-W., Kim, S.-H. & Kim, K.-W. Hypoxia-inducible-factor-HIF—its-p. Exp. Mol. Med. 36, 1–12 (2004).

44 Semenza, G. L. Signal transduction to hypoxia-inducible factor 1. Biochem. Pharmacol. 64, 993–998 (2002).

45 Semenza, G. L. Hydroxylation of HIF-1: oxygen sensing control of oxygen-regulated gene. Physiology 19, 176–182 (2004).

46 Grimes, D. R., Jansen, M., Macauley, R. J., Scott, J. G. & Basanta, D. Evidence for hypoxia increasing the tempo of evolution in glioblastoma. Br. J. Cancer 123, 1562–1569 (2020).

47 Kamura, T. et al. Activation of HIF1α ubiquitination by a reconstituted von Hippel-Lindau (VHL) tumor suppressor complex. Proc. Natl. Acad. Sci. USA 97, 10430–10435 (2000).

48 Amelio, I. & Melino, G. The p53 family and the hypoxia-inducible factors (HIFs): determinants of cancer progression. Trends Biochem. Sci. 40, 425–434 (2015).

49 Sethi, N., et al. Mutant p53 induces a hypoxia transcriptional program in gastric and esophageal adenocarcinoma. JCI Insight 4, e128439 (2019).

50 Amelio, I. et al. p53 mutants cooperate with HIF-1 in transcriptional regulation of extracellular matrix components to promote tumor progression. Proc. Natl. Acad. Sci. U. S. A. 115, E10869–E10878 (2018).

51 Khromova, N.V., Kopnin, P.B., Stepanova, E.V., Agapova, L.S. & Kopnin, B.P. p53 hot-spot mutants increase tumor vascularization via ROS-mediated activation of the HIF1/VEGF-A pathway. Cancer Lett. 276, 143–151 (2009).

52 Raichle, M.E. & Gusnard, D.A. Appraising the brain’s energy budget. Proc. Natl Acad. Sci. USA 99, 10237–10239 (2002).

53 Clark, D.D. & Sokoloff, L. Basic neurochemistry: molecular, cellular, and medical aspects. In: Siegel, G.J., Agranoff, B.W., Albers, R.W., Fisher, S.K. & Uhler, M.D. (eds) Basic Neurochemistry, 7th edn, 637–670 (Lippincott, Philadelphia, 1999).

54 Mattiesen, W.R. et al. Increased neurogenesis after hypoxic-ischemic encephalopathy in humans is age related. Acta Neuropathol. 117, 525–534 (2009).

