## Supplementary material for "Gain-of-function mutant p53 regulates long-noncoding RNA LINC00643 to modulate HIF1α in glioblastoma": Supplimentary file

**Supplementary Figures, Legends, Materials and Methods, Tables**

**Supplementary Figure Legends**

**Figure S1: ChIP confirmation of GOF-MUT-p53 binding lncRNAs in U373 cells.**

**Figure S2: LINC00643 is highly expressed in normal human brain and downregulated in GBM and correlated with survival.** S2A) S2B) NCBI and GTEx databases show high expression of LINC00643 in normal brain. S2C) LINC00643 is downregulated in TCGA GBM tumors. S2D) LINC00643 is downregulated in LGG and GBM tumor in CGGA database. S2E) GTEx transcript database shows junction expression of Exon1-Exon3 of LINC00643 is highly expressed at multiple sites of normal human brain.

**Figure S3. Verification LINC00643 induction by doxycycline treatment in Tet-inducible GBM cells and xenografts by qRT-PCR.** S3A) Validation of Tet-inducible stable U251, U87 and G34 cells that express LINC00643 after Doxycycline treatment for 24 hours. S3B) Expression of LINC00643 after Doxycycline treatment in GBM xenografts. S3C) Validation doxycycline induction of different Exons of LINC00643 in Tet-inducible GBM cells.

**Figure S4: RCAS-Tva mouse model and H&E staining of U87 xenografts.** S4A) Schematic figure of the RCAS-Tva mouse model. S4B) RCAS plasmid construction for overexpression of human LINC00643 and its Exons, and mouse homologues of LINC00643 and Exon3. S4C) H&E staining of U87 xenograft that overexpress LINC00643, Exon3 or control. Since the maximum plasmid size for transduction efficiency (ef) is about 6.4kb and the pRCAS vector has a capacity of about 3.5 kb, we inserted a partial version of LINC00643 that includes Exon1, 2, 3 and conserved region of Exon 4 (Fig. S4B).

**Figure S5 ChIRP-seq to determine LINC00643 binding to DNA.** S5A) Schematic figure of ChIRPseq to uncover lncRNA binding partners. S5B) Probe design and grouping. S5C) Chromatin preparation for U251-LINC00643-expressing stable cells at hypoxic and normoxic growth condition. S5D) ChIRP-seq analysis of binding sites of LINC00643 to DNA under hypoxia and normoxia. S5E) List of top deregulated LINC00643-associated genes under hypoxic and normoxic conditions. S5F) Gene map of the LINC00643 (positive control) and GAPDH (negative control) genes under hypoxic and normoxic conditions, showing differential binding potential for LINC00643 under hypoxic conditions. S5G) Gene set enrichment analysis (GSEA) of low LINC00643 expression samples leads to enrichment of hallmark hypoxia genes S5H) LINC00643 expression correlates with hypoxia pathway gene expression in TCGA GBM Samples S5I) GTEx Portal dot plot shows that HIF1α expression is low in normal brain compared to other GTex tissue types.

**Figure S6. Knockdown of GOF-MUT-p53 reduced HIF1α expression in U373 GBM cells.**

#### Figure S1

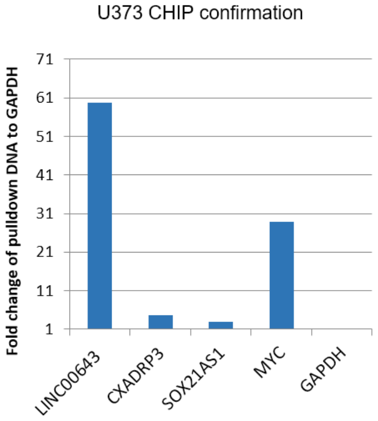


#### Figure S2

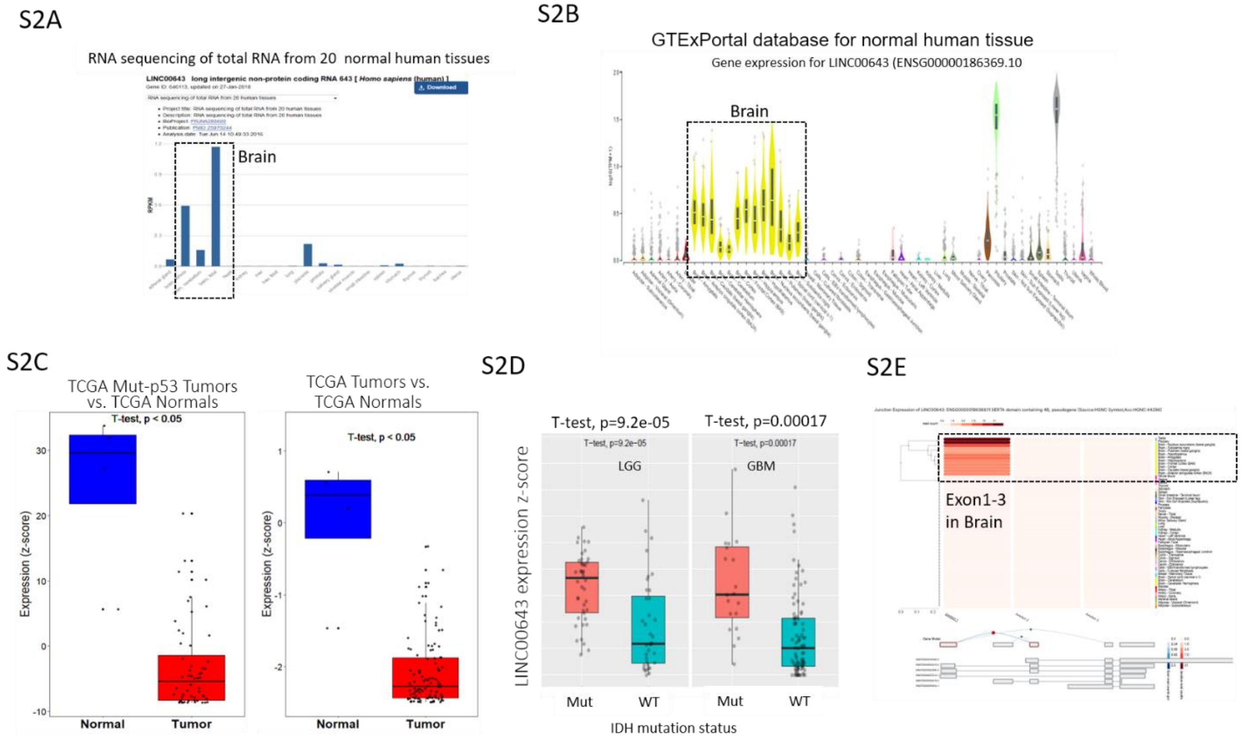

### Figure S3

S3A

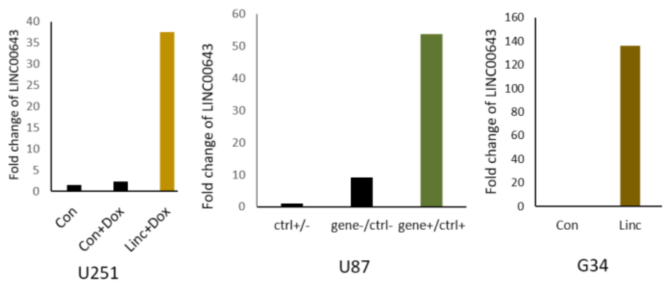


S3B

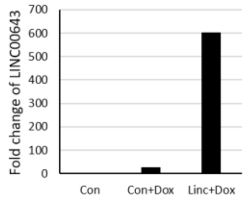


S3C

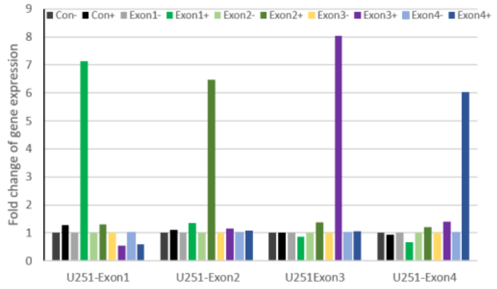

### Figure S4

S4A

RCAS-Tva mouse model

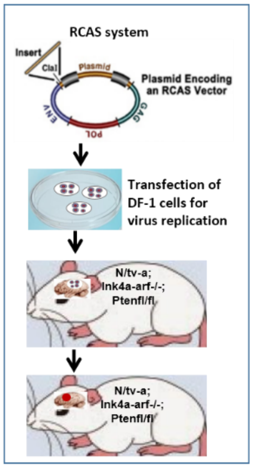


| Group | Injection | Co-injection |
|---|---|---|
| Control | pRCAS-Con | pRCAS-PDGFb pRCAS-Cre |
| LINC00643 | pRCAS-LINC00643 | |
| Exon3 | pRCAS-Exon3 | |

S4B

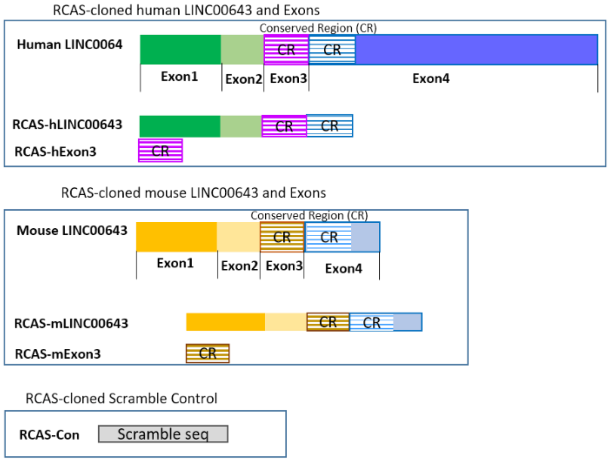


S4C

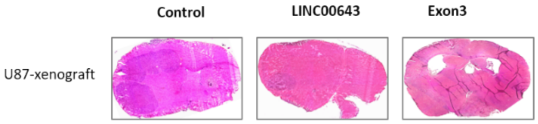


S4D

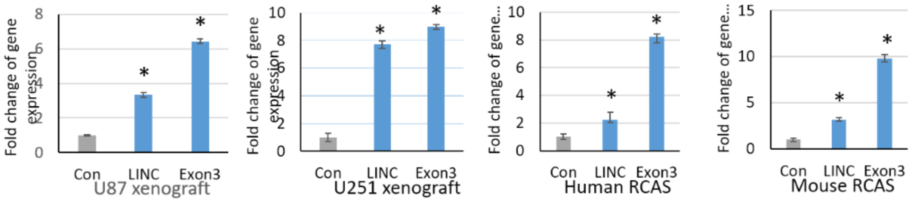

**Figure S5**

S5A

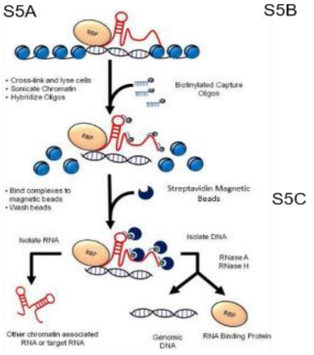


Chromatin isolation by RNA purification (CHIRP) and deep sequencing (CHIRP-seq) to determine the LINC00643 binding partner

S5B

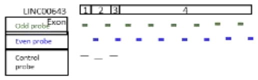


S5C

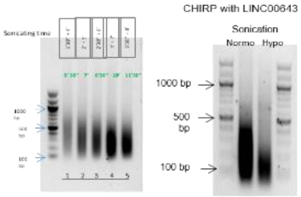


S5D

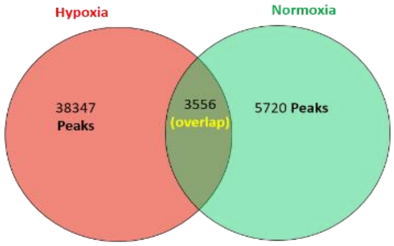


S5E

Hypoxia

| Gene | MeanExpression | FoldChange | padj |
|---|---|---|---|
| HOXB9 | 62.005 | 55.441 | 0.000 |
| TGFBI | 25531.773 | 33.005 | 0.000 |
| VEGFA | 40996.278 | 8.150 | 0.000 |
| HIF1A | 26300.304 | 3.028 | 0.000 |
| HIF1A-AS2 | 9651.283 | 3.209 | 0.000 |
| PFKFB3 | 18107.295 | -1.199 | 0.503 |
| BCL2 | 2129.494 | -1.339 | 0.169 |
| PKLR | 148.496 | -1.697 | 0.028 |
| PRKCA | 10336.822 | -2.071 | 0.000 |
| KIF5A | 17525.733 | -9.242 | 0.000 |

Normoxia

| Gene | MeanExpression | FoldChange | padj |
|---|---|---|---|
| B4GALNT2 | 27.48067 | 2.502252 | 7.22E-05 |
| GCNT2 | 925.522 | 2.141201 | 0.009205 |
| PDK1 | 2632.548 | 1.844468 | 0.038544 |
| EXT1 | 2647.455 | 1.276323 | 0.21246 |
| TPST2 | 3474.39 | 1.239601 | 0.44319 |
| TPBG | 1061.658 | 1.084807 | 0.925443 |
| SCARB1 | 6816.868 | -1.07934 | 0.736635 |
| RBPJ | 9009.867 | -1.19422 | 0.362867 |
| ZNF292 | 3059.249 | -1.525 | 0.02845 |
| NDRG1 | 33872.88 | -1.61054 | 0.334494 |
| MXI1 | 7051.239 | -1.78319 | 0.000563 |
| PDK3 | 3452.611 | -2.98103 | 3.60E-06 |

S5F

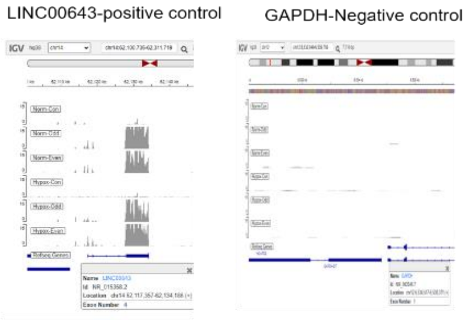


S5G

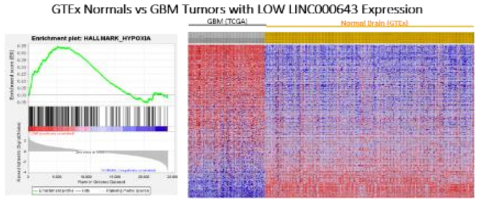


S5H

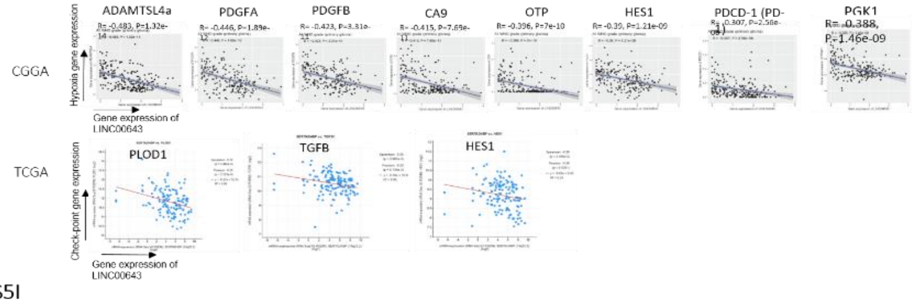


S5I

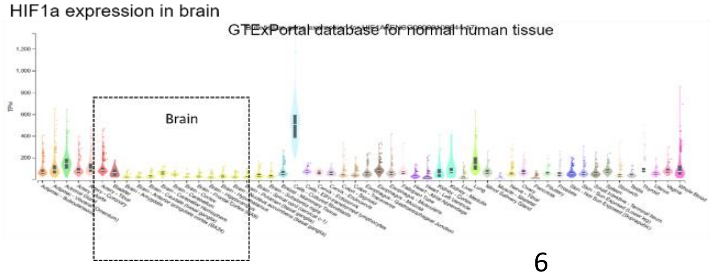

**Figure S6**

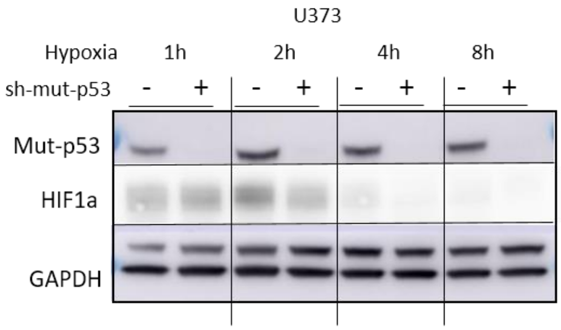


**Supplementary Materials and Methods**

Reagents

All cell culture media and Pen/Strep were from Life Technologies (Grand Island, NY, USA). Fetal Bovine Serum was from GemininBio (West Sacramento, CA). Plasmocin for mycoplasma elimination in cell culture were from InvivoGen. All primers for real-time PCR and cloning were from Life Technologies (Grand Island, NY, USA). Human recombinant EGF and FGF for GSC culture supplements were from R&D Systems (Minneapolis, MN, USA). N2, B27 and L-Glutamine were from Life Technologies (Grand Island, NY, USA). Oligofectamine transfection reagent and Lipofectamine RNAimax were from Life Technologies (Grand Island, NY, USA). X-tremeGENE transfection reagent was from Sigma (St. Louis, MO). The Dual Luciferase kit was from Roche (Pleasanton, CA, USA). The RNeasy Extraction kit were from Qiagen (Chatsworth, CA, USA). iScript cDNA synthesis kit was from BioRad (Philadelphia, PA). All probes for CHIRPseq and scFish were from Integrated DNA Technology (San Jose, CA, USA). All primers for PCR are from Life Technologies (Grand Island, NY, USA).

Vectors and cloning

pTetOne Vector (Takara, Mountain View, CA) was used for construction of Tet-induciable LINC00643 full length and individual Exons. Lenti-vector pCDH-EF1-MCS-BGH-PGK-GFP-T2A-Puro (System Biosciences, Palo Alto, CA) was used for reduced length of LINC00643 and Exons for GSC expression due to the extreme low efficacy of plasmid transfection. pENTR1A (Promega, Madison, WI, USA) was used for LINC00643 and Exon3 cloning followed by gateway cloning into pRCAS.

Transfection and infection

To assess LINC00643 effects on GSC sphere formation, 293TN cells were co-transfected with 2.5 µg plasmid mixture of lentiviral plasmid encoding LINC00643 or Exons, together with psPAX2 and pMD2.G using Xtrem DNA transfection Reagent. After 72 h, viruses were harvested by passing through a 0.45 mm filter. Collected lentivirus was used directly to infect GSC cells with the addition of 8 mg/ml polybrene (Sigma, St. Louis, MO) or stored in − 80 °C. Infected cells were selected with flow sorting by GFP.

Quantitative RT-PCR

Total RNA was extracted from normal human astrocyte, GBM cells, GSCs, and tissues using the miScript RNA extraction kit (Qiagen, Chatsworth, CA, USA)), according to the instructions of the manufacturer. Each RNA sample was reverse transcribed using iScript cDNA synthesis Kit (Bio-Rad, Cat# 1708891). and quantitative PCR analysis was performed utilizing PowerUp SYBR Master Mix (Thermo Fisher, Hillsboro, OR), using the 7500 Real-time PCR System (Applied Biosystems, Carlsbad, CA, USA). LncRNA expression was assessed by fold change calculated using the delta-delta Ct method, with normal brain, normal human astrocytes or untreated cells as a control sample to normalize the fold change.

Subcellular fraction extraction

For nuclear and cytoplasmic RNA separation, $1 \times 10^6$ cells were collected and used for nuclear and cytoplasmic NRA extraction using using PARIS™ kit (Thermo Fisher, Hillsboro, OR) followed by cDNA synthesis and qRT-PCR mentioned above.

CHIP confirmation

CHIP confirmation was performed using purified pulldown DNA fragment by p53 antibody (PCR extraction kit, Qiagen, Chatsworth, CA, USA) as DNA template followed by qPCR for promoter region of LINC00643, CXADRP3, SOX21-AS1, c-Myc and GAPDH using the following primers, FW5'- cccccaaccactgaagagta -3' and RV5' - cacataggcagcgaagtcaa-3' for human LINC00643, FW5'- ttggaggaccattgccttac -3' and RV5'- gcgagggttctcttgttcag -3' for CXADRP3, FW5'- ctcctcctcggtcgttctct -3' and RV5'- ctttggcacttggcagcac -3' for SOX21-AS1, FW5'- ggaaaacgggaatggttttt -3' and RV5'- gatatgcggtccctactcca -3' for Myc, FW5'- gctctctcccatcccttctc -3' and RV5'- ttgcctgtccttcctagctc -3' for GAPDH intron.

Immunoblotting

Cells were lysed in RIPA buffer (50 mM Tris-HCl pH 8.0, 150 mM NaCl, 5 mM EDTA, 0.1% SDS, and 1X protease and phosphatase inhibitor) and loaded (25 µg/lane) and separated onto NuPage Tris Gel, (Thermo Fisher, Hillsboro, OR), and transferred to a nitrocellulose membrane. The membrane was blocked with 5% skim milk and then incubated overnight with gentil rocking at $4^0$C with primary antibodies targeting specific proteins, p53 and HIF1α (1:1000, Cell Signaling, Danvers, MA), GAPDH (1:1000, Santa Cruz Biotechnology, Dallas, TX), After primary antibody incubation, the membrane was probed with horseradish peroxidase-conjugated secondary antibodies at room temperature for 1 hour. Detection was achieved using a pico-sensitive ECL reagent (Thermo Scientific, Waltham, MA), and images were captured using a Kodak Min-R Mammography or Amersham 600 imager.

Cell proliferation assays

U87 and U251 stable cells ($1x10^4$ - $5X10^4$) were seeded and treated with Doxycycline (1 µg/ml) for 48 hours. The cells were collected and counted at various time points for seven days and growth curves were established

Cell invasion assay

To assess LINC00643 effects on GBM cell invasion, we performed transwell invasion assay (Ref SS *66*). In brief, transwell invasion chambers were coated with 150 µl of a 125 µg/ml collagen IV solution. U251 stable cells that overexpress LINC00643, Exon1, Exon2, Exon3, Exon4 or scrambled control were counted, and 1 x $10^5$ cells were re-suspended in 300 µl of 1% serum-containing medium. The cells were added to each transwell chamber and then the chambers were transferred to wells in 24 well plate containing 750 µl of 10% serum-containing medium in the bottom chamber. This created a gradient serum that stimulates cell invasion through the collagen IV-coated membrane. Following 8 hours of incubation, the invaded cells were stained with crystal violet, and five random fields were captured and counted using ImageJ software.

Cell migration assay

To evaluate LINC00643 effect on cell migration, we conducted the cell migration (or scratch) assay. The U251 stable cells($3x10^5$) were seeded in 6 well plate to near confluence, followed by creating a linear scratch across the cell monolayer using a sterile pipette tip. After washing away debris, fresh medium was added, and the cells are incubated 24 hours. Images of the wound are captured. The extent of wound closure and cells migrating in the wound gap is analyzed by measuring the gap area using image analysis software (ImageJ).

GBM stem cell sphere formation assay

GBM stem cells G34 were transduced with lentivirus encoding human LINC00643, Exon3 or control for 48 hour and sort by flow with GFP positive cells. The cells were cultured for 72 hour and were dissociated into single cells with dissociation buffer (EDTA 1 mM, BSA 0.5% in PBS. 1000 single cells were transferred to 24-well plates which pre-coated with poly-ornithine solution (Sigma, St. Louis, MO) and cultured in a neurobasal growth medium at 37°C for 7 days. Neurospheres containing > 30 cells were quantified and represented in a bar graph.

Immunohistochemistry

Tumor bearing mice were euthanized after Ketamine anesthesia with cervical dislocation followed by transcardial perfusion with 25 ml of PBS followed with 15 ml of 4% paraformaldehyde (PFA) (Thermo Fisher, Hillsboro, OR). Brains were dissected out and fixed in 4% PFA for 48 hours at 4C. Mouse brains were then soaked in 30% sucrose (Fisher Scientific) for 5-7 days. Brains were embedded in Tissue plus OCT (Thermo Fisher, Hillsboro, OR) then sectioned in the coronal plane at 12 µM using a cryostat (Thermo Cryostar NX50, Thermo Fisher, Hillsboro, OR). For immunohistochemistry, coronal sections were incubated in a blocking solution (5% normal donkey serum, 0.3% Triton X-100 in PBS) at room temperature for 20 minutes. Rabbit anti-Ki67 (Abcam, Cambridge, UK), mouse anti-NDRG1 and mouse anti-Glut1 (Cell Signaling, Danvers, MA), were diluted in primary diluting solution (5% normal donkey serum, 0.3% Triton X-100 in PBS) and incubated overnight at 4 $^{o}$C. Sections were washed four times in PBT and incubated with Alexa 488 donkey anti-Rabbit or Alexa 647 donkey anti-mouse at 1:250 (Thermo Fisher, Hillsboro, OR) in secondary diluting solution at room temperature for 4 hours. Sections were washed with PBT and mounted with ProLong Gold Antifade Mountant with DAPI (Thermo Fisher, Hillsboro, OR).

**Supplementary Tables**

Table S1. Primers for cloning, qPCR, CHIP and CHIRP

| **Oligo name** | **Forward primer** | **Reverse primer** |
|---|---|---|
| Cloning primers | | |
| Tetone LINC00643 | atatatgaattcGAACAGCTGCGCGCCCGCCTAG | aataatggatccTTTCACCAGAAATAAAATTTTATTAG |
| Tetone exon1 | atatatgaattcGAACAGCTGCGCGCCCGCCTAG | aataatggatccCTTGTGAGCTGGGTCCACTCACCA |

| | | |
|---|---|---|
| Tetone exon2 | atatatgaattcATCTCAGGTAACCATTCATG | aataatggatccCTGGTAATGCTGAATCAATCC |
| Tetone exon3 | atatatgaattcCTTATCCAACAAAAGTATTAAC | aataatggatccTTTTGGAAAATACCCATGG |
| Tetone exon4 | atatatacgcgtCACCTGATATTACCTGAG | aataatggatccTTTCACCAGAAATAAAATTTTATTAG |
| pCDH LINC00643 | aataatttcgaaGAACAGCTGCGCGCCCGC | atatatgcggccgcTTTCACCAGAAATAAAATTTTATTAG |
| qPCR primers | | |
| LINC00643 | GCCTGAGGCTGTGAGAAATGC | GGCTACTTTCGACGTTGTTAATG |
| Exon1 | GTGAGAGGCGCTTCAGAGAC | CGCATGCACCTGTTAGGTAG |
| Exon2 | GCCTGAGGCTGTGAGAAATGC | GTAATGCTGAATCAATCCGAAA |
| Exon3 | AACAACGTCGAAAGTAGCCT | TGCAAGTTCTGTCCTCAGGT |
| Exon4 | GGGCTACACTAGCAGCCACT | TCAGCATGCATCTCATCTTTG |
| CXADRP3 | ATTACATCCTCTGCCTATAAAAATCC | GCACTCAAGATAGAGAAAATC |
| SOX21AS1 | GCAGGAGAGTTAAGGAAAAC | GACTCTCCACTCGCCTAAAC |
| hHIF1α | TGCTTGGTGCTGATTTGTGA | GGTCAGATGATCAGAGTCCA |
| hHIF2α | GTGCTCCCACGGCCTGTA | TTGTCACACCTATGGCATATCACA |
| VEGFA | CACACAGGATGGCTTGAAGA | AGGGCAGAATCATCACGAAG |
| PDK1 | GGAGGTCTCAACACGAGGTC | GTTCATGTCACGCTGGGTAA |
| GLUT1 | TGGACCCATGTCTGGTTGTA | ATGGAGCCCAGCAGCAA |
| JMJD1a | TCAGGTGACTTTCGTTCAGC | CACCGACGTTACCAAGAAGG |
| b-Actin | CCTGTACGCCAACACAGTGC | ATACTCCTGCTT GCTGATCC |
| NDRG1 | AATGCAGAGTAACGTGGAAGTGGTC | TGGTCGCTCAATCTCCAGGTC |
| ANKRD37 | AGGCCAGCACATAGGAGAGA | TTTCTTGCGCTTTCGTTTTT |
| PDGFB | CCATTCCCGAGGAGCTTTATG | GGTCATGTTCAGGTCCAACTC |
| 18sr | CGGCTACCACATCCAAGGAA | GCTGGAATTACCGCGGCT |
| GAPDH | CCATGGAGAAGGCTGGGG | ACCAAAGTTGTCATGGATGACC |
| CHIP primer | | |
| **LINC00643** | cccccaaccactgaagagta | cacataggcagcgaagtcaa |
| CXADRP3 | ttggaggaccattgccttac | gcgagggttctcttgttcag |
| SOX21AS1 | ctcctcctcggtcgttctct | ctttggcacttggcagcac |
| cMYC | ggaaaacgggaatggttttt | gatatgcggtccctactcca |
| GAPDH-intron | gctctctcccatcccttctc | ttgcctgtccttcctagctc |
| CHIRP primer | | |
| TERC | CGC TGT TTT TCT CGC TGA CT | GCT CTA GAA TGA ACG GTG GAA |
| GAPDH_RNA | GTC GGA GTC AAC GGA TTT G | TGG GTG GAA TCA TAT TGG AA |
| WNT1 | AGG GCT GGA ATT TCA AAG GT | TTC TCC TCA GGA TGT ACC CG |
| GAPDH-DNA | GGC TCC CAC CTT TCT CAT CC | GGC CAT CCA CAG TCT GG |
| HIF1α _1 | GAACAGAGAGCCCAGCAGAG | CCTGAGGTGGAGGCGGGTTC |
| HIF1α _2 | AAAGGAAGGGCTTGCTGC | GAAGGGATTTCGGTTGCC |
| HIF1α _3 | AGAGGCTCGGAGCCGG | CTCGTGAGACTAGAGAGAAGCG |
| HIF1α _N | TGCTCATCAGTTGCCACTTC | AAAACATTGCGACCACCTTC |

Table S2. smRNA-FISH probe sequence

| Oligo | Sequence (5'-> 3') with Reporter Dye Quasar 570 (green) |
|---|---|
| LincFL_Probe-1 | gagatcttgtgagctgggtc |
| LincFL_Probe-2 | tcaggctcatgaatggttac |
| LincFL_Probe-3 | acagcgagcatttctcacag |
| LincFL_Probe-4 | aaaatctggtgagagttccc |
| LincFL_Probe-5 | cttttgttggataagctggt |
| LincFL_Probe-6 | aagtaggctactttcgacgt |
| LincFL_Probe-7 | aggtgttttggaaaataccc |
| LincFL_Probe-8 | aagttctgtcctcaggtaat |
| LincFL_Probe-9 | gcttctccagagaaagtttg |
| LincFL_Probe-10 | tctgggtcttcaagaaacct |
| LincFL_Probe-11 | cacagaccttcttaagtagg |
| LincFL_Probe-12 | gtattcatagctctctttct |
| LincFL_Probe-13 | cagtcttataacagggagct |
| LincFL_Probe-14 | taccataaacttcagccgtt |
| LincFL_Probe-15 | gagactgagaacagcattcc |
| LincFL_Probe-16 | gatacgaatgcagctcttca |
| LincFL_Probe-17 | ctccgaagcataaggcacaa |
| LincFL_Probe-18 | tagcccatttcataaatggc |
| LincFL_Probe-19 | agaattttgctccaagtggc |
| LincFL_Probe-20 | actggtaatgagtgccttat |
| LincFL_Probe-21 | gcatgcatctcatctttgat |
| LincFL_Probe-22 | taaccattgctctctccaat |
| LincFL_Probe-23 | gattctaaggactgctgagt |
| LincFL_Probe-24 | agcactaattgaccgcttta |
| LincFL_Probe-25 | taattgtttaacgtaggccc |
| LincFL_Probe-26 | tactttccagtaagagtggc |
| LincFL_Probe-27 | cagggacatggcattatcaa |
| LincFL_Probe-28 | attaaatgtgttccacccctt |
| LincFL_Probe-29 | ccttcctcatatcatttaca |
| LincFL_Probe-30 | ggcacaatgcttgacaagtc |
| LincFL_Probe-31 | ccatctcgaattgctactaa |
| LincFL_Probe-32 | tcagtttggagtaatctggt |
| LincFL_Probe-33 | ctccccgcaaaatagtacaa |
| LincFL_Probe-34 | tagataagcccatctgcaag |
| LincFL_Probe-35 | aaaggcccaccacaaatttt |
| LincFL_Probe-36 | gcagccttaaatgtactctt |
| LincFL_Probe-37 | aaacttcaccaccaaaggca |
| LincFL_Probe-38 | tctgacatctcactcaaacc |
| LincFL_Probe-39 | atgatatgctccataggaca |
| LincFL_Probe-40 | aaatacacctttggcaccta |

| | |
|---|---|
| LincFL_Probe-41 | agtcttcctaacactcatct |
| LincFL_Probe-42 | gcaatccttactttagattc |
| LincFL_Probe-43 | acaatgttcttctatgtcct |
| LincFL_Probe-44 | caaatcattccaaacccctt |
| LincFL_Probe-45 | cccttgattaccattattat |
| LincFL_Probe-46 | gattcctaaaggtctgtagt |
| LincFL_Probe-47 | tagattgttttccaaggcat |
| LincFL_Probe-48 | actttgttgacaagtcacca |

Table S3. CHIRP probe sequence

| Oligo | Sequence (5'-> 3') with Biotin |
|---|---|
| LincProbe_2 | gagatcttgtgagctgggtc/3BioTEG/ |
| LincProbe_4 | acagcgagcatttctcacag/3BioTEG/ |
| LincProbe_5 | aaaatctggtgagagttccc/3BioTEG/ |
| LincProbe_9 | gcttctccagagaaagtttg/3BioTEG/ |
| LincProbe_10 | tctgggtcttcaagaaacct/3BioTEG/ |
| LincProbe_11 | cacagaccttcttaagtagg/3BioTEG/ |
| LincProbe_13 | gatacgaatgcagctcttca/3BioTEG/ |
| LincProbe_14 | ctccgaagcataaggcacaa/3BioTEG/ |
| LincProbe_16 | gattctaaggactgctgagt/3BioTEG/ |
| LincProbe_18 | cagggacatggcattatcaa/3BioTEG/ |
| LincProbe_19 | gcacttcagaaagaaccagc/3BioTEG/ |
| LincProbe_22 | catctgcaaggctatttcgt/3BioTEG/ |
| LincProbe_23 | aactaaaggcccaccacaaa/3BioTEG/ |
| LincProbe_24 | aaacttcaccaccaaaggca/3BioTEG/ |
| LincProbe_25 | tctgacatctcactcaaacc/3BioTEG/ |
| LincProbe_26 | gcatccaactagggaagata/3BioTEG/ |
| LincProbe_27 | caggctaattagggaaccaa/3BioTEG/ |
| LincProbe_28 | acttacagacacggtcctaa/3BioTEG/ |
| LincProbe_29 | cagcaaagcaagcctgtatt/3BioTEG/ |
| LincProbe_32 | gcaaacagtagatatgccga/3BioTEG/ |
| LincProbe_34 | tgggtcaggaaaactttcct/3BioTEG/ |
| LincProbe_35 | gcccacaataaatatgccac/3BioTEG/ |
| LincProbe_36 | cagggatgggctaaggtcag/3BioTEG/ |
| LincProbe_37 | accacacgcacaagatatgc/3BioTEG/ |
| LincProbe_39 | gtgtgggtgaagaatagagc/3BioTEG/ |
| LincProbe_41 | cagacatgaacacagagtct/3BioTEG/ |
| LincProbe_43 | tgccaagcaagaagttgcta/3BioTEG/ |
| LincProbe_44 | tgttgtagaacaccatctgc/3BioTEG/ |
| LacZ_1 | ccagtgaatccgtaatcatg/3BioTEG/ |
| LacZ_2 | tcacgacgttgtaaaacgac/3BioTEG/ |
| LacZ_3 | attaagttgggtaacgccag/3BioTEG/ |
| LacZ_4 | aggttacgttggtgtagatg/3BioTEG/ |

| | |
|---|---|
| LacZ_5 | aatgtgagcgagtaacaacc/3BioTEG/ |
| LacZ_6 | gtagccagctttcatcaaca/3BioTEG/ |
| LacZ_7 | aataattcgcgtctggcctt/3BioTEG/ |
| LacZ_8 | agatgaaacgccgagttaac/3BioTEG/ |
| LacZ_9 | aattcagacggcaaacgact/3BioTEG/ |
| LacZ_10 | tttctccggcgcgtaaaaat/3BioTEG/ |
| LacZ_11 | atcttccagataactgccgt/3BioTEG/ |
| LacZ_12 | aacgagacgtcacggaaaat/3BioTEG/ |
| LacZ_13 | gctgatttgtgtagtcggtt/3BioTEG/ |
| LacZ_14 | ttaaagcgagtggcaacatg/3BioTEG/ |
| LacZ_15 | aactgttacccgtaggtagt/3BioTEG/ |
| LacZ_16 | ataatttcaccgccgaaagg/3BioTEG/ |
| LacZ_17 | tttcgacgttcagacgtagt/3BioTEG/ |
| LacZ_18 | atagagattcgggatttcgg/3BioTEG/ |
| LacZ_19 | accattttcaatccgcacct/3BioTEG/ |
| TERC_1 | aggcccaccctccgcaaccc/3BioTEG/ |
| TERC_2 | aaaaatggccaccaccccctc/3BioTEG/ |
| TERC_3 | ggcgcctacgcccttctcaa/3BioTEG/ |
| TERC_4 | acagcgcgcggggagcaaaa/3BioTEG/ |
| TERC_5 | ccgctgaaagtcagcgagaa/3BioTEG/ |
| TERC_6 | cggcaggccgaggcttttcc/3BioTEG/ |
| TERC_7 | ctctagaatgaacggtggaa/3BioTEG/ |
| TERC_8 | ccagcagctgacatttttttg/3BioTEG/ |
| TERC_9 | aggtccccgggaggggcgaa/3BioTEG/ |
| TERC_10 | ttcgggggctgggcaggcga/3BioTEG/ |
| TERC_11 | cgaccgcggcctccaggcgg/3BioTEG/ |
| TERC_12 | tgcctccggagaagccccgg/3BioTEG/ |
| TERC_13 | aactcttcgcggtggcagtg/3BioTEG/ |
| TERC_14 | agacccgcggctgacagagc/3BioTEG/ |
| TERC_15 | tgaacctcgccctcgccccc/3BioTEG/ |
| TERC_16 | tcttcctgcggcctgaaagg/3BioTEG/ |
| TERC_17 | gcgcggggactcgctccgtt/3BioTEG/ |
| TERC_18 | tcccacagctcagggaatcg/3BioTEG/ |
| TERC_19 | tgagccgagtcctgggtgca/3BioTEG/ |

Table S4. Human LINC00643 cDNA sequence (6636bp)

>NR_015358.2
gaacagctgcgcgcccgcctagcgctgcgaacccggagtgagaggcgctt
cagagactggagggacggacctgcaggagccagggctgcccgccgagctg
cagttgccgctactgccgctgtcagcgcccgactgaagagtgacagccgc
aaaagcaggagtgagcaccgaaccccgcgcttctacctaacaggtgcatg
cgccgatctggtgagtggacccagctcacaagatctcaggtaaccattca
tgagcctgaggctgtgagaaatgctcgctgtgcaaaggggaactctcacc
agattttctgaaactgttctagaaatacacttttcggattgattcagcat
taccagcttatccaacaaaagtattaacatatacattaacaacgtcgaaa

gtagcctactttaagagaaaatatgcagaagaagaagatttacatcaaga
ttaccatgggtattttccaaaacacctgatattacctgaggacagaactt
gcattctcaaactttctctggagaagctcaggtttcttgaagacccagaa
acctacttaagaaggtctgtgttaattaacaatttgctgaggaagataca
tctagagacggagaaagagagctatgaatactttaaagaagctccctgtt
ataagactgcatattctgatacaagaaaacggctgaagtttatggtacag
gaatgctgttctcagtctctctattatgaagagctgcattcgtatcatat
tgtgccttatgcttcggagaatgccatttatgaaatgggctacactagca
gccacttggagcaaaattctcagttgcttatttataaaatgaattaaagt
aactgttaagtccattgatactttgtttctaaatgcttgaactgtttata
gtgaagatgttaatataaggcactcattaccagtgcacttgatatcaaag
atgagatgcatgctgaactgattggagagagcaatggttattactcagca
gtccttagaatcatcaaattgaatagaaaatatgttcacatatgataaag
cggtcaattagtgctttagaagcaaagatttttttaaacataagatgtg
taggaaggggcctacgttaaacaattacacagatgaaaaattttctttta
tgccttgatttccaaattctgccactcttactggaaagtaatgaaagtaa
cagattaaattgataatgccatgtccctgaggatgaagggtggaacacat
ttaattttctgtaaatgatatgaggaagggctggttctttctgaagtgct
ttgacttgtcaagcattgtgccaaataattgacaccattctccttttacc
attagtagcaattcgagatggggcaaagacactgcaactaaactgctgaa
tgtaacaaccagattactccaaactgatccattgtactattttgcgggga
gggcttagacttttctctttaaaagtgaattttaacgaaatagccttgca
gatgggcttatctataagttctacaggttggttaaaatttgtggtgggcc
tttagttagccgtttacattttatttgttttttttataggctgtgaagc
tcaggaaaagggtaggcaggtaaaatttgtttttaatgagttgatacttt
ttagaaaccttatctgcaaaaattaatgagaacaacaaattccacatgat
gttatttcttcatctagctaagagtacatttaaggctgcctggatgaaac
tttgcactgatttttgtataatagttgatcattcactatttgacaaattg
tttcagaatgcctttggtggtgaagtttagataagtccctatgttttgca
attttacagataattatttaacaagttggatagattcatgaaattgtttg
aagtggttcatactgttctttcaggtttgagtgagatgtcagatttcagt
gtctgaagttttgtcctatggagcatatcatttaaagaattgtcattaac
tgataaaacagattggaaaaactaggtgccaaaggtgtatttaagaaaat
accaaagactcaaagatgagtgttaggaagactatgataatgaagataag
aatctaaagtaaggattgcataggacatagaagaacattgttaaaggggt
ttggaatgatttgcttactaatgtactttcagaacactttatacataata
atggtaatcaagggcactgctttaaatgaagatactacagacctttagga
atcatatgccttggaaaacaatctatgtatggtgacttgtcaacaaagta
tcttccctagttggatgccaacttgtaaccaagaagactttcttctattc
aagagaaatttggttttagttttgatttaatacttggtccctctaattaa
tcactggggtgattaacatgaattctgttggtataaagagtattggttcc
ctaattagcctgcattgttgaagaccacatagatgtattactgataagac
tatgaggctcatgtttaattactcaatcaaaattaggaagaaatgctttg
gaagaaactaaaaaattacaatttaggttaatttttaaaggcaatttagt
atacatgatgagctagctttgaaatacagatttaaatgtttaaaaattaa
tttgagggcaagatttggtaccaaagtttcaaaaattttatttcactgct
tataaaatactttatcagaacacttattttaaaaaataacatcttatttg
aaaaactccaaatttgatgggatagttacaccatcatacaattttaaagg

tagaattagctttctaatactttaaaatattttctgaaatgaacatttca
cttggtttaggaccgtgtctgtaagtaaaagacaccaatgctaatagata
tgacttacaaaaatgccacatatcgaaattgattgaggattgaaatatat
atactttggcattgagaagagaggaagagaaaaaaataaaaaaagtgtga
cattgtaatatttattgtatttaataaatcttgaagaaagaactcttgag
caatgaaccagtgacagatataaaagcctgttttatttgatgacatcatt
aactgagttttctcatctatacattcatacatcgagaaggaactgtctta
aaaaccagtgtttctttttttttttaagttctagcgtacatgtgcagaat
gtgcaggtttgttatataggtatacacgtgccatggtggtttgctgcacc
catcaacctgtcatctacattaggtatttctcctaatgctatccctcccc
agtgatatttgttaatatgcttttttaaaaaagtgtctgccttgaaactg
gtaattaaaaaatatctcctagggcatttaattatttttcccaataaaca
actcatagataacgggttgactaatgttatctttatttagaaaatttgta
tgtaaactgtgtaaacttgtctgagttaaccaatagactaccggggtttt
agcatggtaaactgatatacagggagacccaaatacaggcttgctttgct
gactacccagcgtgctttattacaggagatgcaaaaggtggaagaccagt
tacaccttttttttttttttttaaaacctggacatcctttctctggtga
caagagccatccttatggtaaggaaggtgagatagaaaatggagaacctt
tggtaatgttgatctttcttgtgggtgtccacctagcctaaaagccaagt
gaagaagaacataaaaaagcagaagaggaaaaatgaagaaaagaggaaaa
agagggtggggccagagaaataaagagtaggattagtaagtgaaagaaaa
agttgctttgttgtgtggggggggtgttcttgcttgctatactcaatttt
gctttcccgtgtctgctgtacacaaaacacctgatctctgcaatgtattg
ctcctttctttcattcacctgtgatgcataagactagattattttcggca
tatctactgtttgcaaagtgttactactgaaaaatatccctgaaactgag
ctctttgggtggataagcaaaggaaaaatagaaaataattaaggtaaggg
aaaggctaaaggataagcctgtgtataaatgggaaatggataagctcaaa
tgcattatctggtttcaatgtaacacccaagatttaacaaactcagtgct
agaagacttgaaaataagtgtaatttaccaccatctattgagcagctatt
atgagccaggcactgtgctagggctggggatacataagtgaataatgcac
agtcccagaactcagattatttggttttgttttaccaaatccaaatgcag
tacctgcatttctcttttccaaactgagatggctatcaaacatgtctttc
agaaagtgtttgcaggtgagaagatgcgcaaggtgaaggaaagttttcct
gacccagatcttagaaggaaaggagaggatacattttgctttgtggcata
tttattgtgggcaaaaagctactattgcctaagggaagtacggctgacct
tagcccatccctggggcatatcttgtgcgtgtggtggggagacaaatcag
gtagggaacaattccttctcgccttacctctctagcttccatgttctttt
atggaacaaatcagattaatactaatgttaaggagagctttaaaggagaa
agagaatcaataaatcacagcctgaaagttgtgtatgttgtgtgcaagct
cagaggggcagtcttcttcaatttgccttgtgctggtgaattgcttgaat
gaacttcggtatttcttaacaccaggtactggagcccaccttctttctct
ccctctggtttcctctttaaatcacagcctgaccccagtctttatagtcc
attgtaagtggaagttatagctctattcttcacccacaccttgctcccta
tcattgatacttagaagaaagtaacaatttgcagtactggctgaactcct
ttgggaaagtttctggagtgtatcaaataagaattcatcatagtaacatg
gtcgttactggctgaacaaaattctttttgagactattgtacttagtcat
taaataattgtttactaaggcaattttcatgtttctggaattcagtgtaa
tagttaacagctgtatatgtctcacaaaagaaactacttaggttggaaac

aatggaaggttgtgtataattaattcaatcaggtcatgaatatttatgta
acatatggcattttaatttatatgttcccattctcatacttcattactat
acagcagcaacaagataaatttcaggtttttgttttttattaagtggg
ccatgtctaaaagttgtcacattcctggttgaatattatggacaaaattt
ccccattaaagtagttttgtctttctcaaggattatcctttagggttggg
tggattaaaaacattacattagtgcttcttgagcatacaagtcactaggg
atcttgtgaaaatacagattccttttagtaggtttgggatgaggaatgaa
ggtcttcatctctcaaatctcccaggtgatgtggatgctgccagtccatc
gaccacactttgagttgggagattctacatcttttgagaaatatccacac
tgaagcctatactcttaaactttcaaagactctgtgttcatgtctgtgtt
ctgcaagaatttttcttttaagaaataaactgcataaagtaaaatcaga
aaaccataacactggttttccaaatttgccacaaatactgtaatactctg
tagagtaaaatgcaaagattattcctgttacaagttttctctgtatcaag
tgcaggaaaggaacatgggtagagtcatgtaccattcttatcagtcagga
gatgacacgtggtaaatttctcttcttgattttcctcttgattatactca
cataagggagctccatttggtaacaaagatgaaattctgttcacagttaa
caagaatttagcaacttcttgcttggcaaaatctgagacaaccttacaaa
aacatcctacattaaattcagaattttgggtagctgcataagctgaagat
tatggaaaacctgagctgaaaatggcacctggatctgtaacttcttgtct
tgaactctttttgagctttattctgtgagagatcttcccctacagtgat
tttttctgtttctcctcagtcgctggggtctcagtaaggggtggaggatt
ggtgtaaatgagacagtcacataaattgtctaatttgagcatgccaagtg
atttttgtcagcctcttttggtcataaaattttggtatagctattgtgaa
atatagtgtcataaatttgtcataagccattaatgaaggaagagaagcag
aaatttatttctgtgggaatgcactcaaatatcaagcagatggtgttcta
caacatttatttgggaaaatgtgtatctgttacataatctgaaatatgtc
tttttcacatttaaaaatatttgggtcatgatttagagtttttattggat
tgtttttaaactgagaggaagaagaaaggtaattgtattttaaaacatt
tgacatgttactaataaaattttatttctggtgaaa

Table S5. Mouse LINC00643 cDNA sequence (1058bp)

>NR_030734.1
gttctgagccggagcgagaggcgcttcagagaaggagggaccgacctgca
ggagccggccgccccgccgagctgcagtcgctgccgccgccgctgtcgct
gtcgctgtcagaacccgtccgaagagtgacaccttcgcgaaaaaacagga
gctgtgaaacatttgcacacgcagtcttcggatcccaggtccccattcat
gagcccgaggctgtgagacatgctccatgtgccaaggggagttctcacca
catcttccgaggctgtcctagacatacactcttcggtctgagtcagcatt
accagacagtgctgctgctacctcgagaaatcgctggccatcagtgtccc
tggaagtccctaagaaagccttgcagaatctgcagcttgttgggaatctg
aaactgccaataaaatcaatattgtgcctggagtagatgataaatctgaa
gaaaaagattactccattgattcagagctaaacaccgacactgaactctg
ctgttatagagcatatcatacagaaagcttatccaagagaagtcctatca
tattcatcaacaacgtcgaaagtagcctactttaagagaaaatatgcaga
agaagaagatctacatcaagattcccatgggtattttccaaaacacctga

tattaccggaagacaggacttgtatcctcaaactttccctagagaagctc
aggtttcttgaagacccagaaacctacctaagaaggtctgtgttaattaa
caatttgctgaggaagatacacctagagacggagaaagagagctatgaat
acttcaaggaagctccctgctataagactgcttattctgatacgagaaaa
cggctgaagtttatggtacaggagtgctgttcccagtctctctactatga
agagctgcactcctaccacattgtgccttacacttcggagagtaccatct
atgggatgggctacactggtgaccacttggagcaaaattctcagttgctc
atttataaaatgaattaaaaagtaactgttaagttcattgaaaaaaaaaa
aaaaaaaa
